# *Tombusvirus* p19 captures RNase III-cleaved double-stranded RNAs formed by overlapping sense and antisense transcripts in *E. coli*

**DOI:** 10.1101/510842

**Authors:** Linfeng Huang, Padraig Deighan, Jingmin Jin, Yingxue Li, Elaine Lee, Shirley S. Mo, Heather Hoover, Sahar Abubucker, Nancy Finkel, Larry McReynolds, Ann Hochschild, Judy Lieberman

**Affiliations:** Program in Cellular and Molecular Medicine, Boston Children’s Hospital, Boston, Massachusetts, USA; Department of Pediatrics, Harvard Medical School, Boston, Massachusetts, USA; Department of Biomedical Sciences, Jockey Club College of Veterinary Medicine and Life Sciences, City University of Hong Kong, Hong Kong SAR, China; Biotechnology and Health Centre, City University of Hong Kong Shenzhen Research Institute, 8 Yuexing 1st Road, Shenzhen Hi-Tech Industrial Park, Nanshan District, Shenzhen, China; Department of Microbiology and Immunobiology, Harvard Medical School, Boston, Massachusetts, USA; Department of Biology, Emmanuel College, Boston, Massachusetts, USA; Division of RNA Biology, New England Biolabs, Ipswich, Massachusetts, USA; Novartis Institutes for Biomedical Research, Cambridge, Massachusetts, USA

**Author notes:** Correspondence should be addressed to L.H. or J.L.

**Keywords:** RNase III, Antisense RNA, p19, Small RNA, dsRNA

## Abstract

Antisense transcription is widespread in bacteria. By base pairing with overlapping sense RNAs, antisense RNAs (asRNA) can form long double-stranded RNAs (dsRNA), which are cleaved by RNase III, a dsRNA endoribonuclease. Ectopic expression of plant tombusvirus p19 in *E. coli* stabilizes ~21 bp dsRNA RNase III decay intermediates, which enabled us to characterize otherwise highly unstable asRNA by deep sequencing of p19-captured dsRNA and total RNA. dsRNA formed at most bacterial genes in the bacterial chromosome and in a plasmid. The most abundant dsRNA clusters were mostly formed by divergent transcription of sense and antisense transcripts overlapping at their 5’-ends. The most abundant clusters included small RNAs, such as *ryeA/ryeB*, 4 toxin-antitoxin genes, and 3 tRNAs, and some longer coding genes, including *rsd* and *cspD*. The sense and antisense transcripts in abundant dsRNA clusters were more plentiful and had longer half-lives in RNase III mutant strains, suggesting that formation of dsRNAs promoted RNA decay at these loci. However, widespread changes in protein levels did not occur in RNase III mutant bacteria. Nonetheless, some proteins involved in antioxidant responses and glycolysis changed reproducibly. dsRNAs accumulated in bacterial cells lacking RNase III, increasing in stationary phase, and correlated with increased cell death in RNase III mutant bacteria in late stationary phase. The physiological importance of widespread antisense transcription in bacteria remains unclear but it may become important during environmental stress. Ectopic expression of p19 is a sensitive method for identifying antisense transcripts and RNase III cleavage sites in bacteria.

## Introduction

Endogenous antisense RNAs (asRNA) are products of DNA-dependent RNA polymerase initiated from antisense promoters that overlap at least partially with any coding or functional RNA (sense RNA). The overlapping regions of sense and antisense RNAs are fully complementary so they have the potential to form perfectly matched double-stranded RNAs (dsRNA). Usually asRNA are much less abundant than the corresponding sense RNA. Next generation sequencing has identified many new species of asRNA (1, 2), but their biological significance is not well understood.

Examples of RNA regulation of gene expression in bacteria have been described that involve a variety of small non-coding RNAs and asRNA (3–6). asRNA and RNase III regulate gene expression of plasmids and toxins. The *E. coli* ColE1 plasmid replication origin encodes two non-coding RNAs – RNA II, which serves as a DNA replication primer, and RNA I, which is shorter and fully complementary to the 5′ portion of RNA II (7–10). RNA I inhibits plasmid replication by binding to the RNA II plasmid replication primer. Other well-known asRNA-regulated systems are the Type I Toxin-Antitoxin (TA) genes (11). In Type I TA systems, a small RNA gene lies opposite to, but overlapping with, a gene encoding a toxic peptide. The small asRNA inhibits the expression of the toxin by at least partially base pairing with the toxin RNA. Examples of asRNA-mediated toxin regulation systems include *hok/sok* in the R1 plasmid (12, 13) and *ldrD/rdlD* in the *E. coli* genome (14). RNase III, an exonuclease that cleaves dsRNAs (15) to generate 5’-phosphate and 3’-hydroxyl termini, leaving a characteristic 3’ 2 nucleotide (nt) overhang (16, 17), regulates both the plasmid replication system^10^ and the Type I TA systems (18, 19). Exhaustive digestion of dsRNAs by RNase III produces small dsRNAs of about 14 bp.

Bacterial genomes produce many asRNAs from protein coding genes. Using a whole-genome tiling microarray, the Church group discovered that a large percentage of the *E. coli* genome is transcribed in both directions (20). Multiple groups subsequently used deep sequencing to study the transcriptome of bacterial genomes (6). Lasa et al. found a significant increase in the number of antisense reads within the short (<50 nt) RNA deep sequencing reads in *Staphylococcus aureus* (2). Their findings suggested that asRNA are widely transcribed across the genome of Gram-positive bacteria but are degraded with sense RNAs into small RNAs <50 nt by RNase III. Lioliou et al. used a catalytically inactive RNase III mutant to pull down RNase III-bound RNAs and identified RNase III-bound asRNA in 44% of annotated genes in *S. aureus* (21). More recently, deep sequencing of immunoprecipitated dsRNAs in an *RNase III* deficient strain (*rnc* mutant) revealed that RNase III degrades pervasive sense and antisense RNA pairs in *E. coli* (22).

Despite the consensus that asRNA are ubiquitous in bacterial genomes, the biological functions and physiological significance of asRNA are not well understood. There are only a few examples of asRNA regulating protein coding genes (23). One study suggested that asRNA are mainly transcriptional noise arising from spurious promoters (24). In contrast, two operons overlapping in their 5’ regions were shown to antagonize each other’s expression in *Listeria monocytogenes* (25). Whether widespread asRNA are ubiquitous gene regulators or mostly transcriptional noise and what is the role of RNase III in asRNA gene regulation remain to be investigated in *E. coli*.

The tombusvirus p19 protein captures siRNAs (~21 nucleotide small dsRNAs) to defend against the antiviral effects of RNA interference in plants (26, 27). We previously found (28) that ectopic expression of p19 in *E. coli* captures ~21 nucleotide small dsRNAs generated from overlapping exogenous sense and antisense transcripts. These small RNA duplexes, which are apparently intermediary degradation products of RNase III, were termed pro-siRNAs (**pro**karyotic short interfering RNAs). Precipitation of p19 in bacterial cells co-expressing *p19* together with ~500 nt sense and antisense sequences or a similarly sized sense-antisense stem-loop of an exogenous gene enabled us to isolate and purify pro-siRNAs that specifically and efficiently knocked down the exogenous gene when transfected into mammalian cells (28–30). pro-siRNAs mapped to multiple sequences in the target gene. In bacteria expressing p19, but no exogenous sequences, ~21 nucleotide dsRNAs were also captured (referred as ‘p19-captured dsRNAs’). These short dsRNAs were greatly reduced in the absence of p19 or in RNase III-deficient bacteria expressing p19. We hypothesized that these short dsRNAs represent p19-stabilized RNase III decay intermediates of overlapping endogenous sense and antisense transcripts that might provide a useful method for characterizing labile endogenous dsRNAs.

## Results

### Plasmid-directed synthesis of p19 uncovered plasmid encoded dsRNAs

Two methods for expressing p19 proteins in bacteria were designed (Fig. 1a). The first method uses a pcDNA3.1 plasmid (pcDNA3.1-p19-FLAG), previously engineered by us (28) to express p19, driven by the CMV promoter. To characterize the dsRNAs captured by p19 pull-down, we compared RNAs isolated after overnight culture from cell lysates of WT (MG1693) and RNase III-deficient *rnc-38* in the MG1693 background (SK7622) (31), transformed with pcDNA3.1-p19-FLAG. In *rnc-38*, insertion of a kanamycin resistance gene within a 40-bp fragment in the *rnc* gene abrogates RNase activity (31). dsRNAs bound to p19 were isolated using affinity chromatography, cloned and deep sequenced. Sequencing reads were mainly 21-22 nt long from WT *E. coli* and were reduced ~10-fold in the RNase III mutant strain (Fig. 1b, c), consistent with our previous finding that pro-siRNAs are produced by RNase III (28). The aligned reads in WT *E. coli* mapped to both the *E. coli* genome and plasmid, but most of the aligned reads (78%) mapped to the plasmid (Fig. 1c). The plasmid reads were unevenly distributed across the entire plasmid, but were concentrated in ‘hot spots’ (Fig. 1d), as previously found in cells expressing exogenous hairpin RNAs (28). The hot spots contained unequal levels of sense and antisense reads, as previously found for exogenous sequences, where the differences in abundance of sense and antisense reads were shown to likely be due to cloning bias (28).

**Figure 1.**
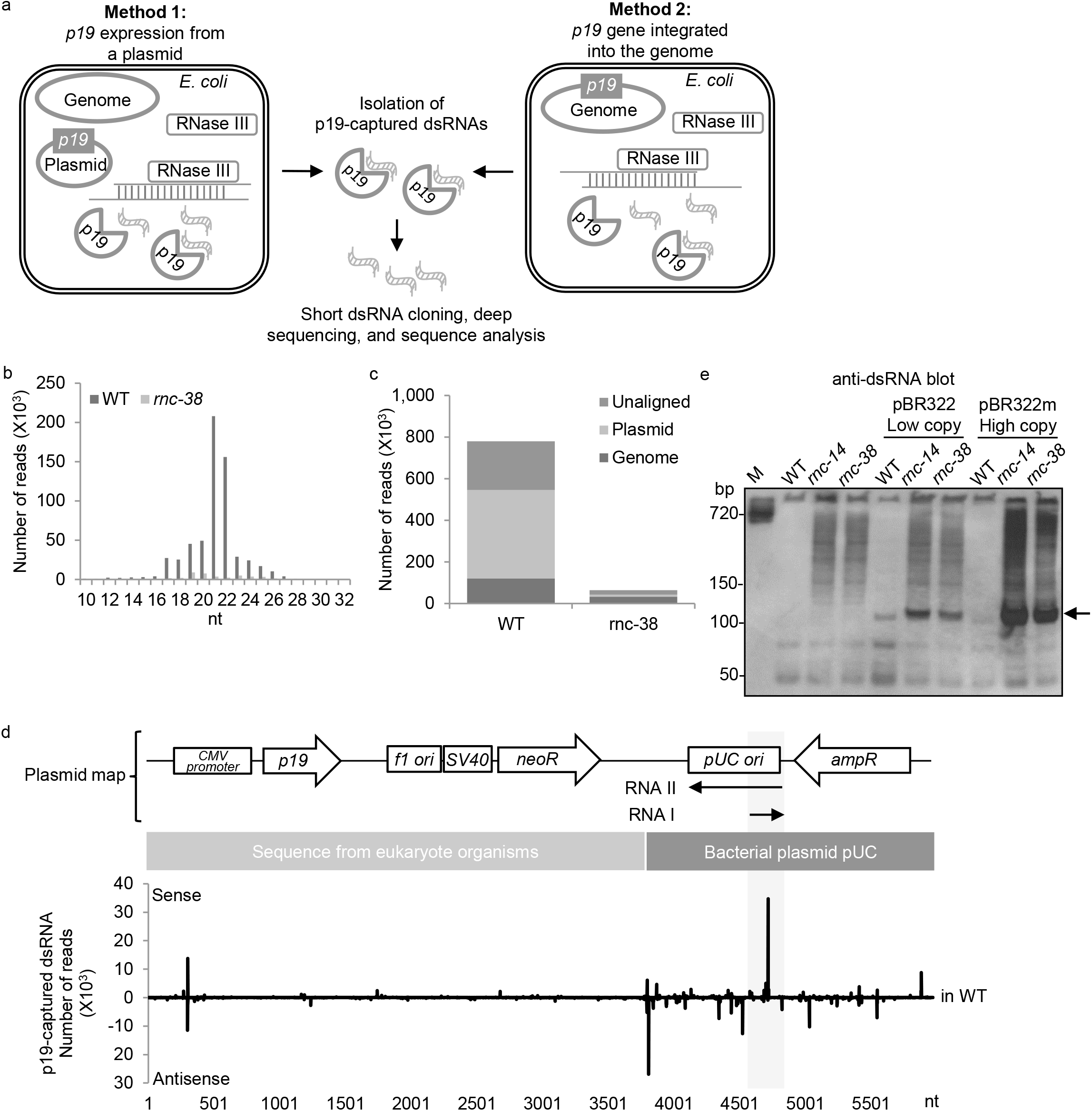
dsRNAs are generated from a plasmid. a. Schematic of two methods used to isolate p19-captured dsRNAs in this study. b. Length distribution of sequencing reads of short dsRNAs isolated from *E. coli* WT (MG1693) and *rnc-38* mutant (SK7622) strains transfected with p19 expressing pcDNA3.1-p19-FLAG plasmid. c. Summary of deep sequencing read alignments. d. Top: map and sequence features of pcDNA3.1-p19-FLAG plasmid; Bottom: plot of p19-captured dsRNA distribution from the plasmid in *E. coli* MG1693 strain. e. dsRNA immunoblot of *E. coli* total RNA probed with J2 antibody. Gel stained with SYBR-Gold before blotting is shown in Supplementary Fig. 1a. M: a dsRNA of 720 bp.

The pcDNA3.1-p19-FLAG plasmid is comprised of a pUC bacterial plasmid backbone, but includes additional sequences supporting functions in eukaryotic cells since pcDNA3.1 was designed for use in mammalian cells (Fig. 1d). p19-captured dsRNA hot spots were most abundant in the sequences of bacterial plasmid origin and distributed along it, suggesting that the bacterial plasmid produces multiple overlapping sense and antisense RNAs, consistent with a recent study (32). By contrast non-bacterial sequences were largely devoid of dsRNAs, except for dsRNAs observed within the CMV promoter region, which drives *p19* expression. We speculate that the other non-bacterial sequences might not be adapted to initiate transcription in bacteria and thus produce fewer overlapping transcripts and fewer dsRNAs. The origin of replication of pcDNA3.1 is derived from the pUC plasmid, which is known to produce a sense RNA I transcript, which promotes replication, and an antisense RNA II transcript, which inhibits replication (33). The overlapping region of RNA I and RNA II contained a dsRNA hot spot, consistent with the idea that dsRNAs originate from overlapping transcripts (Fig. 1d).

Our results suggested that RNase III is responsible for degrading dsRNAs produced from both bacterial genome and plasmid. dsRNAs made from the plasmid were more abundant than those made from the genome. To confirm RNase III-dependence, we isolated total RNA from WT and *rnc* mutant *E. coli*, which were transformed or not with a low or high copy number version of pBR322 (pBR322 wild type or mutant pBR322 high copy number). *rnc-14* (34) and *rnc-38* (31) are both RNase III deficient by previous studies and by our RNA sequencing data. Equal amounts of RNA from each condition were separated by PAGE, transferred to nitrocellulose membranes and detected with J2 antibody, which recognizes dsRNA (Fig. 1e, Supplementary Fig. 1a). dsRNAs of various sizes were detected but were much more abundant in *rnc* mutant strains. The amount of total dsRNA was higher in plasmid-containing bacteria, particularly those bearing the high copy variant. A prominent band of ~100 bp (indicated by arrow in Fig. 1e) was only detected in plasmid-containing bacteria. Thus, consistent with dsRNA sequencing, plasmids produced abundant dsRNAs, which exceeded the amount of genomic dsRNAs (Fig. 1c). There was no noticeable plasmid copy number difference between the WT and *rnc* mutant strains, ruling out the possibility that the increase in dsRNA seen in the mutant strains was due to an increase in plasmid DNA and suggesting that RNase III does not regulate plasmid copy number. The relative abundance of plasmid-derived dsRNA sequences in transformed bacteria might be because pcDNA3.1 is a high copy number plasmid that generates more transcripts than the single copy bacterial genome.

### p19-captured dsRNA hot spots are caused by sequence bias of RNase III and are not strongly skewed by p19

The hot spot pattern, observed in the plasmid p19-captured dsRNAs, raised a concern that p19 capture might have sequence bias. To examine whether p19 capture introduces any substantial sequence bias that could skew the distribution of short dsRNAs, two exogenous gene long dsRNAs were generated by annealing T7 RNA polymerase-transcribed complementary sense and antisense sequences of *LMNA* or *eGFP*, gel purified and incubated *in vitro* with *E. coli* RNase III (NEB) (E1-3) or human Dicer (Genlantis) (E4). Two reaction conditions for *E. coli* RNase III were compared – one that contained Mg^2+^ (E1) and one that contained Mn^2+^ (E2) as the divalent cation. p19 pulldown was also used to capture ~21 nt dsRNAs produced in the Mn^2+^ reaction (E3). The *in vitro* RNase III and Dicer cleavage products, with and without p19 capture, were cloned and sequenced (Supplementary Fig. 2a). Because of the known cloning bias of different strands of dsRNAs (28), the sense and antisense reads after *in vitro* digestion were combined to plot the total reads along each sequence. As expected (35), RNase III digestion in the presence of Mg^2+^ mostly produced small RNAs of ~14 bp, while digestion in the presence of Mn^2+^ or Dicer digestion produced predominantly ~21 bp dsRNAs (Supplementary Fig. 2b).

**Table 1.** Top 15 p19-captured dsRNA clusters overlapping with small RNAs.

| Cluster No | Genome coordinates | RPM in S2 | Small RNA | Functions | Lybecker dsRNA | Conway overlapping transcripts |
| --- | --- | --- | --- | --- | --- | --- |
| C2-155 | 1,921,124-1,921,555 | 19,505.4 | *ryeA, ryeB* | *cis*-acting; overlapping small RNAs | Y | Y |
| C2-220 | 2,812,816-2,812,884 | 9,452.4 | *micA* | *trans*-acting | Y | Y |
| C2-307 | 3,698,166-3,698,199 | 5,552.8 | *rdlD, ldrD* | Type I TA | N | N |
| C2-273 | 3,348,440-3,348,771 | 5,448.9 | *arcZ* | *trans*-acting | Y | Y |
| C2-123 | 1,489,381-1,489,535 | 5,086.0 | *rydC* | *trans*-acting | Y | N |
| C2-134 | 1,620,782-1,620,946 | 4,442.6 | *mgrR* | *trans*-acting | Y | N |
| C2-166 | 2,041,159-2,041,583 | 4,024.7 | *serU* | tRNA | Y | Y |
| C2-170 | 2,151,328-2,151,397 | 2,296.0 | *ibsA, sibA* | Type I TA | Y | N |
| C2-333 | 4,047,984-4,048,014 | 2,202.0 | *spf* | *trans*-acting | Y | N |
| C2-255 | 3,192,754-3,192,892 | 2,030.7 | *ibsD, sibD* | Type I TA | Y | N |
| C2-2 | 16,960-17,328 | 1,930.3 | *mokC, sokC* | Type I TA | N | Y |
| C2-375 | 4,526,006-4,526,367 | 1,923.7 | *ryjB* | *trans*-acting | Y | Y |
| C2-263 | 3,316,137-3,316,391 | 1,887.7 | *metY* | tRNA | N | Y |
| C2-304 | 3,662,902-3,662,980 | 1,765.7 | *gadY* | *trans*-acting | Y | Y |
| C2-232 | 2,945,132-2,945,598 | 1,575.1 | *metZ, metW, metV* | tRNA | N | Y |

RPM: Reads Per Million.

All short RNA products generated *in vitro* by bacterial RNase III in both conditions or human Dicer showed hot spots (Supplementary Fig. 2c). Although the hot spot patterns were all somewhat different, many of the peaks coincided between the samples. While RNase III cuts dsRNA at any position, it was surprising that human Dicer, which supposedly cuts siRNA from the end of dsRNA, also produced many internal short RNA peaks, suggesting that Dicer cleavage has sequence preferences. Short dsRNAs pulled down by p19 from RNase III Mn^2+^ reaction products (E3) displayed a similar distribution pattern as all short dsRNAs generated in the Mn^2+^ reaction (E2). E2 and E3 profiles were highly correlated (Pearson’s correlation coefficient, r=0.74 for *LMNA* sequence, *p*-value<0.0001; r=0.34 for *eGFP* sequence *p*-value<0.0001), but sequences generated under different conditions or by RNase III versus Dicer were less similar, suggesting that RNase III class enzymes have sequence preferences, but p19 capture does not introduce significant sequence bias.

### Features of genomic dsRNAs

Because p19 capture does not introduce substantial sequence bias, we deep sequenced p19-captured dsRNAs to map endogenous labile dsRNAs produced from sense and antisense transcription of the bacterial genome. To focus on genome-encoded dsRNAs, *p19* was integrated into the lambda phage attachment site of the MG1655 *Δlac* genome (36) (Method 2, Fig. 1a) and its expression was driven by an IPTG-inducible *tac* promoter. p19-bound short dsRNAs were isolated after IPTG induction in both exponential (sample ‘S1’) and stationary (‘S2’) phases (Supplementary Fig. 3a), cloned and sequenced using a SOLiD deep sequencer. Approximately 20 million reads that aligned to the *E. coli* genome were obtained from each sample (Supplementary Table 1). dsRNAs were generated from most *E. coli* genes in both samples and the abundance of reads for each gene in the two samples was highly correlated (r=0.846), suggesting that bacterial growth stage does not affect dsRNA production globally (Fig. 2a). The abundance of sense and antisense p19-captured RNA reads from each gene were roughly equal, as expected for dsRNAs (r=0.705, Fig. 2b). In contrast, sense reads were much more abundant than antisense reads in total RNA, analyzed by deep sequencing of total RNA after rRNA depletion (‘RNA-seq’, sequencing reads in Supplementary Table 2) (r=0.059, Fig. 2c). The level of p19-captured dsRNAs varied greatly among *E. coli* genes (Fig. 2a). 83% of genes in S1 and 87% of genes in S2 produced at least 1 dsRNA Read Per Million reads (RPM) (Fig. 2d), indicating that most bacterial genes have at least some partially overlapping antisense transcripts. Comparing the level of p19-captured dsRNA sequencing reads with RNA-seq sense or antisense reads, p19-captured dsRNA reads correlated significantly better with antisense RNA-seq reads for both exponential growth and stationary phase datasets than with sense RNA-seq reads (Supplementary Fig. 3b). Thus, p19-captured dsRNAs are generated widely across the *E. coli* genome and their abundance is related to asRNA transcription.

**Figure 2.**
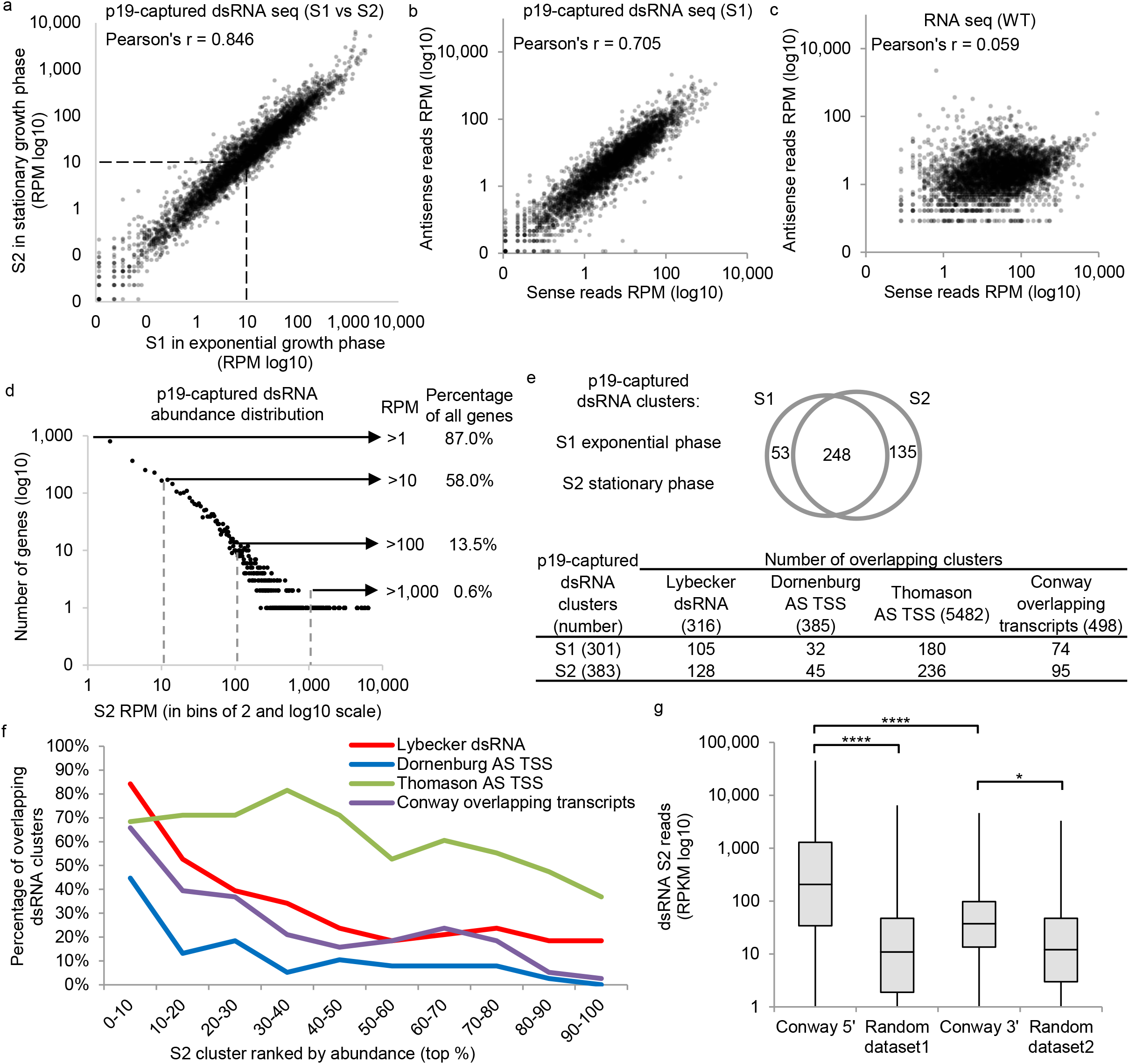
Analysis on *E. coli* genomic p19-captured dsRNA datasets. a. XY plot of p19-captured dsRNA reads, S1 versus S2. b. XY plot for sense and antisense reads of a p19-captured dsRNA deep sequencing dataset (S1). c. XY plot for sense and antisense reads of a total RNA deep sequencing dataset from WT *E. coli*. For a-c, each data point represents an *E. coli* gene annotated by RefSeq. Data are from Supplementary Table 1 (p19-captured dsRNA datasets) & 2 (total RNA datasets). d. Distribution of the abundance of p19-captured dsRNA in all *E. coli* genes. e. Top: Venn diagram shows the number of overlapping clusters between S1 and S2; Bottom: comparing p19-captured dsRNA clusters with dsRNA transcriptome dataset (Lybecker dsRNA), two antisense (AS) transcription start site (TSS) datasets (Dornenburg AS TSS and Thomason AS TSS), and an overlapping transcript dataset (Conway). f. Percentage of overlapping loci according to the abundance of p19-captured dsRNA for S2 clusters. Details about p19-captured dsRNA clusters are in Supplementary Table 3. g. Box plot of dsRNA read density (RPKM in log10 scale) in 5' or 3' overlapping regions of the operons identified by Conway et al. The random datasets have the same length distribution as the experimental datasets. *: *p*-value<0.05; ****: *p*-value<0.0001.

### p19-captured dsRNAs cluster in well-defined genomic loci

To study the dsRNAs that most likely originated from longer dsRNAs formed by overlapping transcription from opposite DNA strands, we focused on clusters of p19-captured dsRNAs with the most sequencing reads. We arbitrarily defined a p19-captured dsRNA cluster as a genomic region that contains at least 2,000 reads within 200 bp. 301 dsRNA clusters were identified in the S1 sample, while 383 were identified in the S2 sample (Fig. 2e, Supplementary Table 3). Most clusters (248) were found in both samples. Because the exponential and stationary phase clusters highly overlapped, we focused on clusters identified in the S2 (stationary phase) dataset in the subsequent analysis.

A previous study (Lybecker et al. (22)) identified sense-antisense paired transcripts, which were stabilized in *rnc-105* mutant *E. coli* (which contain an *rnc* missense mutation encoding a protein with <1% WT RNase III activity (37)), by immunoprecipitation with a dsRNA-specific antibody (J2). 316 dsRNA clusters were identified in Lybecker et al. (22). About a third of the p19-captured dsRNA clusters we identified in either the S1 or the S2 sample overlapped with the Lybecker dsRNA loci (Fig. 2e). These 128 overlapping transcripts (for S2) were concentrated in the clusters with the highest number of reads. The clusters were assigned to 10 groups based on the abundance of reads in each cluster. 84% of clusters in the most abundant group were identified as dsRNA loci by Lybecker et al. (22), suggesting that p19-captured dsRNA clusters are indeed generated from dsRNAs (Fig. 2f). This high degree of concordance suggests that the most abundant clusters are unlikely to be transcriptional noise.

If p19-captured dsRNA clusters are formed by overlapping sense and antisense transcripts, as we hypothesize, they should contain known antisense transcription start sites (asTSS). Two recent studies, Dornenburg et al. (23) and Thomason et al. (38), used deep sequencing to identify asTSS. S2 dsRNA clusters were compared with these asTSS datasets. Thomason et al. identified the largest number of asTSSs (5,482), which were present in more than 50% of dsRNA clusters, while Dornenburg identified 385 asTSSs. When we ranked the S2 clusters by the abundance of reads, the most abundant clusters overlapped more strongly with the predicted asTSSs (Fig. 2f). Within the top 10% most abundant S2 clusters, 45% and 68% contained asTSSs identified by Dornenburg and Thomason, respectively, suggesting that p19-captured dsRNA abundance is a good indicator of the extent of antisense transcription and that p19-captured dsRNA abundance could indicate the extent of overlapping sense and antisense transcripts at a genomic locus.

### p19-captured dsRNA clusters are enriched at the 5’ overlapping regions of operons

Recently Conway et al. used high resolution strand-specific RNA deep sequencing, promoter mapping, and bioinformatics to predict operons (39). The full-length operons they defined included some operons that overlapped at their 5’ ends (divergent operons) or 3’ ends (convergent operons). In total they identified about 500 overlapping transcripts including 89 novel antisense transcripts and 18 coding transcripts that completely overlapped with operons on the opposite strand. 95 of the 383 S2 clusters overlapped with the overlapping operons identified by Conway et al. (Fig. 2e). Again, more abundant p19-captured dsRNA clusters overlapped more with the Conway dataset (Fig. 2f). The read density (RPKM, Reads Per Kilobase Million) of all p19-captured dsRNAs that overlapped with Conway’s divergent operons (129 loci with median RPKM of 206.3) was significantly greater (*p*-value<0.0001) than the read density of p19-captured dsRNAs that overlapped with Conway’s convergent operons (255 loci and median RPKM is 37.3) (Fig. 2g). The RPKM for these overlapping clusters were also significantly greater than randomly generated sets of p19-captured dsRNA clusters of similar size (for divergent operons, median RPKM of random dataset was 11.0; for convergent operons, median RPKM of random dataset was 12.2). These results suggest that RNase III-produced short dsRNAs captured by p19 are more often generated from divergent transcripts.

### Characterization of top dsRNA clusters

To focus on dsRNAs that are more likely to be biologically relevant, we analyzed the more abundant clusters, because most of these were also identified in other studies (Fig. 2e, f). The top 15 p19-captured dsRNA S2 stationary phase clusters involving known small RNA genes, which had 1,575-19,505 RPM, are listed in Table 1 and the top 20 stationary phase p19-captured dsRNA clusters involving only protein coding genes, which had 4,478-17,960 RPM, are listed in Table 2. To understand the features of these more abundant p19-captured dsRNA clusters, we plotted the p19-captured dsRNA seq and RNA-seq reads onto the annotated genome for *E. coli* MG1655 in the UCSC genome browser, together with the published dsRNA and TSS data (Fig. 3, 4 and Supplementary Fig. 4–7). To investigate whether and how RNase III regulates transcripts overlapping with the dsRNA clusters, Northern blots of total RNAs, extracted from WT and *rnc* mutant strains (*rnc-14* or *rnc-38*), were probed for sense and antisense transcripts of some of the abundant p19-captured dsRNA cluster genes. To test whether sense or antisense RNA stability is affected by RNase III, RNA half-lives were also examined for some clusters by comparing Northern blot sense and antisense signals in WT and *rnc* mutant bacteria harvested at various times after adding rifampicin to block *de novo* transcription.

**Figure 3.**
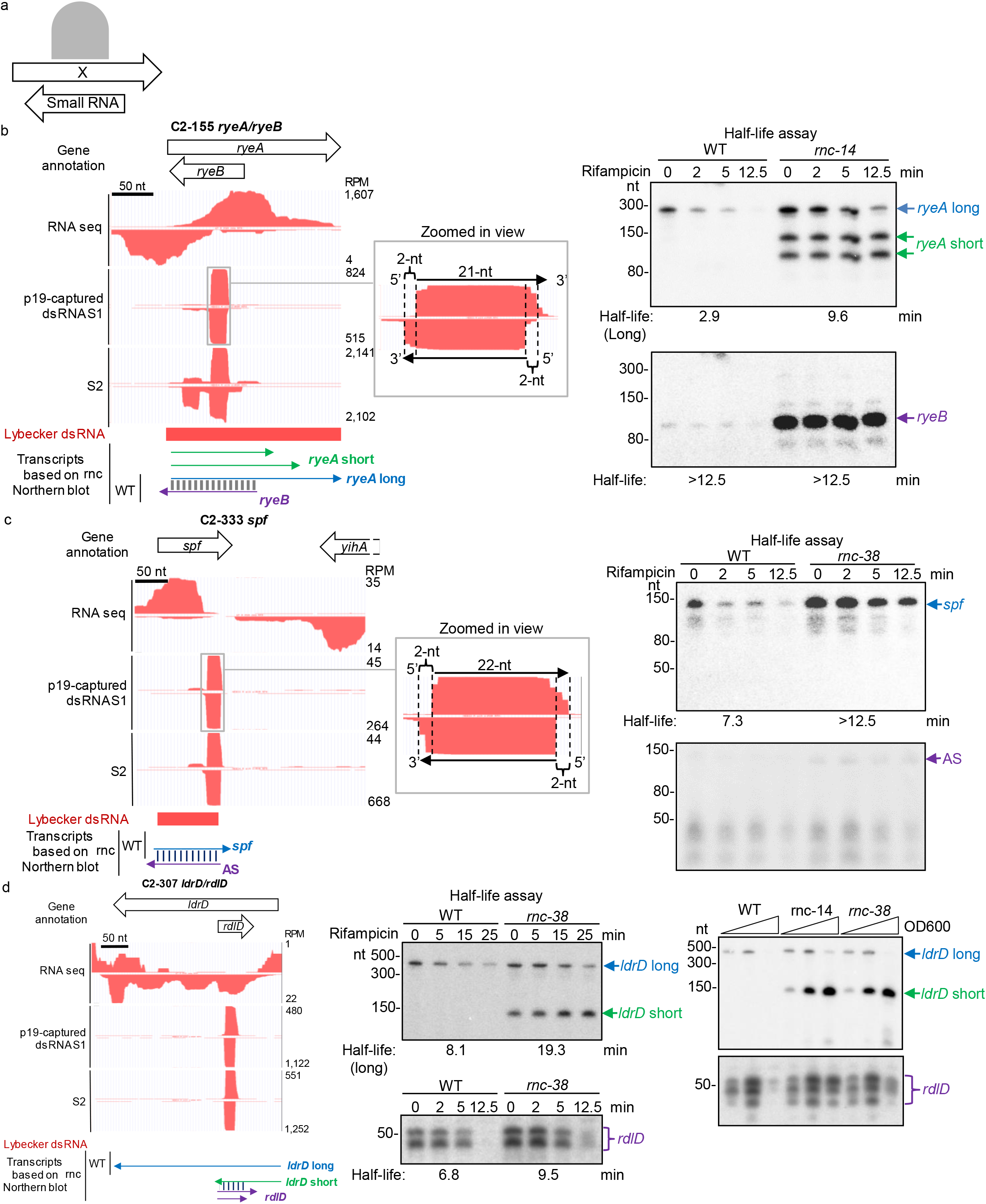
p19-captured dsRNA clusters in small RNAs. a. Schematic showing how antisense transcripts overlap with small RNAs. b. *ryeA/ryeB* locus. Left: p19-captured dsRNA and total RNA sequencing data were plotted in UCSC genome browser and predicted RNA transcripts are shown. Middle: zoomed-in view of the indicated small dsRNA peak. Right: RNA half-life assay. c. *spf* small RNA locus. Same set of data are shown as in b. d. *IdrD/rdlD* locus. Left: genome browser snapshot; Middle: RNA half-life assay; Right: Northern blots of RNA samples isolated during bacterial growth at indicated times. Arrow and bracket indicate putative RNA transcripts and the color scheme matches that in genome browser snapshot.

**Table 2.** Top 20 p19-captured dsRNA clusters (S2) overlapping with protein coding genes.

| Cluster No | Genome coordinates | RPM in S2 | Genes | Type of overlap | Lybecker dsRNA | Conway overlapping transcripts |
| --- | --- | --- | --- | --- | --- | --- |
| C2-324 | 3,963,390-3,964,342 | 17,960.5 | *rhlB, trxA* | 5' | Y | Y |
| C2-349 | 4,194,368-4,195,338 | 17,842.2 | *rsd, nudC* | 5' | Y | Y |
| C2-325 | 3,988,343-3,989,253 | 11,520.6 | *hemC, cyaA* | 5' | Y | Y |
| C2-43 | 437,393-437,579 | 10,668.9 | *yajO, dxs* | New AS | N | Y |
| C2-372 | 4,483,986-4,484,275 | 9,421.8 | *pepA, lptF* | 5' | Y | Y |
| C2-328 | 4,014,014-4,014,166 | 9,113.6 | *ysgA, udp* | 5' | Y | Y |
| C2-116 | 1,386,335-1,386,859 | 8,326.6 | *tyrR, tpx, ycjG* | Full | Y | Y |
| C2-382 | 4,632,313-4,633,370 | 8,181.6 | *rob, creA* | 5' | Y | Y |
| C2-339 | 4,103,836-4,104,050 | 7,910.5 | *cpxR, cpxP* | 5' | Y | Y |
| C2-332 | 4,024,830-4,025,594 | 7,881.9 | *fre, fadA* | 3' | N | N |
| C2-169 | 2,111,741-2,112,327 | 7,686.3 | *galF, wcaM* | Full & New AS | Y | N |
| C2-76 | 918,037-918,309 | 7,674.3 | *ybjX, macA* | 5' | Y | Y |
| C2-355 | 4,254,212-4,254,971 | 7341.3 | *plsB, dgkA* | 5' | Y | Y |
| C2-188 | 2,411,203-2,411,785 | 6,875.4 | *yfbV, ackA* | 5' | Y | Y |
| C2-283 | 3,440,668-3,441,270 | 6,611.1 | *rpmJ, secY* | New AS |  |  |
| C2-383 | 4,638,370-4,638,791 | 6,119.6 | *arcA, yjjY* | Full | Y | Y |
| C2-77 | 921,163-922,104 | 5,810.9 | *macB, cspD* | Full | Y | Y |
| C2-361 | 4,391,754-4,392,109 | 5,634.8 | *yjeS, yjeF* | 5' | Y | Y |
| C2-298 | 3,572,736-3,572,888 | 4,716.9 | *asd, yhgN* | 5' | Y | Y |
| C2-259 | 3,291,446-3,291,566 | 4,477.5 | *yraL, yraM* | 5' | Y | Y |

RPM: Reads Per Million.

### RNase III-regulated small RNAs

Bacterial small RNAs are typically 50-300 nt in length, can code for small peptides or be noncoding, and include important gene regulators (4). 14 of the 301 S1 clusters and 18 of the 383 S2 clusters overlapped with known small RNA genes (Supplementary Table 3). A schematic of dsRNAs overlapping with small RNAs is shown in Fig. 3a. Most of the 15 most abundant dsRNA clusters (11 of 15) were also identified as dsRNAs by Lybecker et al. (22) and 8 of 15 were identified as overlapping transcripts by Conway et al. (39) (Table 1). *ryeA/ryeB* (also known as *sraC/sdsR)*, a locus that may play a role in bacterial responses to environmental stress (40, 41), was the top stationary phase p19-captured dsRNA cluster (19,505 RPM) (Table 1). A previous study showed that *ryeB* (104 nt) regulates the level of *ryeA* (249 nt) in a growth and RNase III-dependent manner (42). The dsRNA sequencing data (in S1) showed one dominant short dsRNA peak of ~21 nt in the overlapping region of *ryeA* and *ryeB* (Fig. 3b, left). Within this hot spot, a zoomed in look at the sequence showed that this 21-nt dsRNA has 3’ 2-nt overhangs at both ends, suggesting that these dsRNAs are *bona fide* RNase III products. Full length and shorter transcripts of both *ryeA* and *ryeB* were more abundant and *ryeA* had a ~3-fold increased half-life in *rnc-14* than WT bacteria (Fig. 3b, right). Two smaller fragments of *ryeA* were detectable and stable only in *rnc* mutants, suggesting that there are alternative labile transcripts or intermediate decay products of *ryeA* that only become detectable when RNase III is not present. Thus, RNase III regulates *ryeA* and *ryeB*, as previously described (42).

Another p19-captured dsRNA cluster in a small RNA is Spot 42 (2,202 RPM), a glucose-regulated small RNA encoded by the *spf* gene, which regulates many metabolic genes in *trans* (43). dsRNA reads were immediately downstream from RNA sequencing reads, suggesting that RNase III cleavage might help form the 3’ end of Spot 42 RNA (Fig. 3c, left). A zoomed in view of the dsRNA hot spot showed that it is a 22-nt dsRNA with 3’ 2-nt overhangs at both ends (Fig. 4c, middle), consistent with production by RNase III. An *spf* asRNA, of the same size as Spot 42 RNA (109 nt), was only detected in the *rnc* mutant (Fig. 4c, right), consistent with results of Lybecker et al. (22). Moreover, *spf* expression and half-life were greater in the *rnc* mutant strain (Fig. 4c, right). We also identified an abundant p19-captured dsRNA cluster (9,452 RPM) for another stress-related small RNA, *micA* (72 nt), which binds to Hfq, is known to be processed by RNase III and inhibit mRNA translation by an antisense mechanism (44). Although the abundance of *micA* was not substantially changed in the *rnc-38* mutant strain, its stability increased (Supplementary Fig. 4a). The antisense transcripts of *micA* were more abundant and several additional short transcripts were detected only in the *rnc* mutant. Thus, RNase III cleaves Spot 42 and *micA* dsRNAs, which regulates their stability. Abundant dsRNAs reads were also identified within *arcZ, rydC, mgrR, ryjB*, and *gadY*, suggesting that there are overlapping antisense transcripts and RNase III cleavage at those loci (Supplementary Fig. 4b).

**Figure 4.**
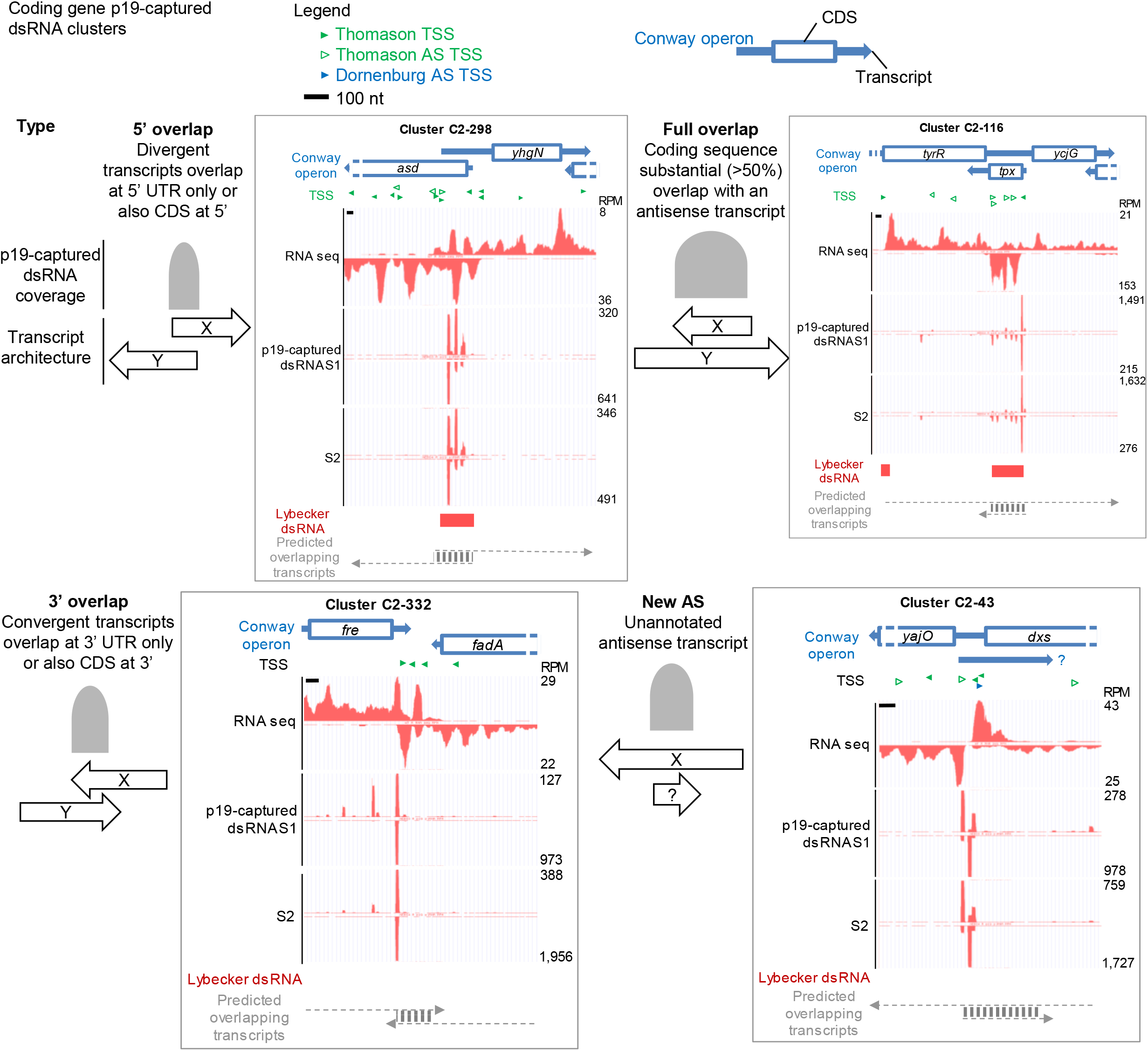
p19-captured dsRNA clusters in coding genes. Schematic shows major types of overlapping sense and antisense transcripts in coding genes. The graph is based on snapshots from the UCSC genome browser. Full length mRNA transcripts were based on annotation by Conway et al. and CDS was marked according to genome annotation in RefSeq. Transcription starts sites (TSS) cloned by Thomason et al. (LB2.0 TSS) are marked by green triangles with empty green triangles highlighting antisense TSSs defined by Thomason et al. Antisense TSS identified by Dornenburg et al. are marked by blue triangles. p19-captured dsRNA sequencing reads (both S1 and S2) and RNA sequencing reads from a WT *E. coli* are plotted in UCSC genome browser. dsRNA identified by Lybecker et al is marked by a red bar. We have arbitrarily predicted potential overlapping transcripts that could give rise to p19-captured dsRNAs and classified all p19-captured dsRNA clusters (Supplementary Table 3).

Three tRNA loci *(metY, serU, metZ/metW/metV)* were also amongst the top 15 small RNA p19-captured dsRNA clusters (Table 1). At these tRNA loci, dsRNA reads were not restricted to the region of the mature tRNAs but also occurred in the surrounding regions (Supplementary Fig. 5). These patterns suggest that longer precursor tRNA transcripts are the source of these dsRNAs.

### RNase III regulates toxin-antitoxin small RNA loci

The Type I toxin-antitoxin (TA) systems involve small non-coding RNA, which can base-pair with a small toxin mRNA to inhibit toxin expression (19). Within the top 15 small RNA p19-captured dsRNA loci were three families of cis-acting TA I loci – *ldr/rdl, mok/sok*, and *ibs/sib. ldrD* encodes for a small toxic peptide and the antisense *rdlD* is an antitoxin small RNA that form a Type I TA system (19). This locus (5,553 RPM) was not identified as forming dsRNA by Lybecker et al. (22). However, a ~21 nt dsRNA peak is present within the overlapping region of *rdlD* and *ldrD* (Fig. 3d). The expression level and half-life of the full-length transcript of *ldrD (ldrD* long) and *rdlD* were both slightly increased in *rnc* mutant strains (Fig. 3d middle). A stable smaller fragment of *ldrD* transcript *(ldrD* short), which accumulated during bacterial growth, was detectable only in the *rnc* mutant (Fig. 3d right). In two other *E. coli* Type I TA loci with overlapping sense and antisense RNAs, *mokC/sokC* and *ibsD/sibD* (45), the stability of the toxin transcripts increased in *rnc* mutants (Supplementary Fig. 6). By contrast, the steady-state level and stability of opposite strand antitoxin small RNAs did not change much in *rnc* mutant strains. In all three Type I TA loci, stable smaller sense RNA fragments were detected only in the *rnc* mutants, which could either be alternative transcripts or degradation products of the full-length transcript (Fig. 3d and Supplementary Fig. 6). Our data suggest that RNase III regulates the expression of toxins by cleaving dsRNAs formed with the toxin mRNA.

Together our data suggest that RNase III regulates the level and/or stability of some mature small RNA transcripts, including both cis-acting (e.g. *ryeA/ryeB* and Type I TA loci) and transacting (e.g. Spot 42 and *micA)* small RNAs.

### Top p19-captured dsRNA clusters in coding genes

The p19-captured dsRNA seq and RNA-seq reads of protein coding gene loci were also mapped onto the annotated genome for *E. coli* MG1655 in the UCSC genome browser, together with the published dsRNA (22) and TSS (38) predictions (Fig. 4). The Conway dataset was used to mark full length transcripts, when available. Based on all the available data, all top p19-captured dsRNA clusters were classified as in previous publications (2, 22, 38) according to whether the sense and antisense transcripts were divergent (5’ overlap), convergent (3’ overlap) or the coding sequence (CDS) overlapped entirely or almost entirely with the predicted antisense transcript (Fig. 4, Table 2, and Supplementary Table 3). dsRNAs were assigned to the latter class if >50% of the CDS was contained in p19-captured reads. A fourth category was defined by abundant dsRNA clusters that did not overlap with previously annotated antisense transcripts. Some clusters contained more than one predicted dsRNA.

Within the top 20 coding gene p19-captured dsRNA clusters, the most common category (13 of 20) involved divergent mRNA transcripts of adjacent genes encoded on opposite strands, which overlap in their 5’ regions. In some cases, dsRNA formed only within the 5’ UTRs but in others included some of the 5’-end of the coding sequence. An example of this category is the C2-298 locus, which contains overlapping 5’ sequences of the *asd* and *yhgN* genes on opposite strands (Fig. 4). p19-captured dsRNAs in this cluster were produced only in the overlapping regions of the RNA transcripts that were predicted by Conway et al. (39) and were supported by our RNA-seq data. At this locus, the dsRNA identified by Lybecker et al. (22) coincided with the region where we sequenced p19-captured dsRNAs. Other examples are shown in Supplementary Fig. 7a. All 13 of the predicted dsRNAs for these abundant divergent clusters overlapped at least partially with dsRNAs pulled down with dsRNA antibody (22), although often they were not identical in position or length.

In the top 20 coding gene clusters, only one cluster arose from convergent transcripts of adjacent genes encoded on opposite strands, which overlap in the 3’ region – the C2-332 locus involving *fre* and *fadA* genes (Fig. 4). At this locus, the 3’UTR of *fadA* mRNA or possibly a transcript initiated from within the 3’ region of the *fadA* gene or 3’ to it (as suggested by previously identified asTSSs and the *fadA* operon mapping by Conway et al. (39)) overlapped with the *fre* transcript. This cluster was not identified by dsRNA pulldown by Lybecker et al. (22).

Another category, full overlap, contains coding genes that overlap substantially with another RNA transcript. 4 of the most abundant 20 clusters fell into this category. An example of this class is the *tpx* gene in C2-116 (Fig. 4). p19-captured dsRNAs were detected across the entire *tpx* CDS. Based on the RNA-seq data, the antisense transcript of *tpx* could come from the 3’ UTR of *tyrR* mRNA, the 5’ UTR of *ycjG* mRNA, a read-through transcript containing both *tyrR* and *ycjG* (as annotated by Conway et al. (39)), or even a new antisense transcript unrelated to *tyrR* and *ycjG*. This cluster and other examples showing a putative overlapping transcript across the entire CDS of *yjjY* in C2-383 (Supplementary Fig. 7b) and *cspD* in C2-77 (Fig. 5a) were also identified by dsRNA pulldown by Lybecker et al. (22).

**Figure 5.**
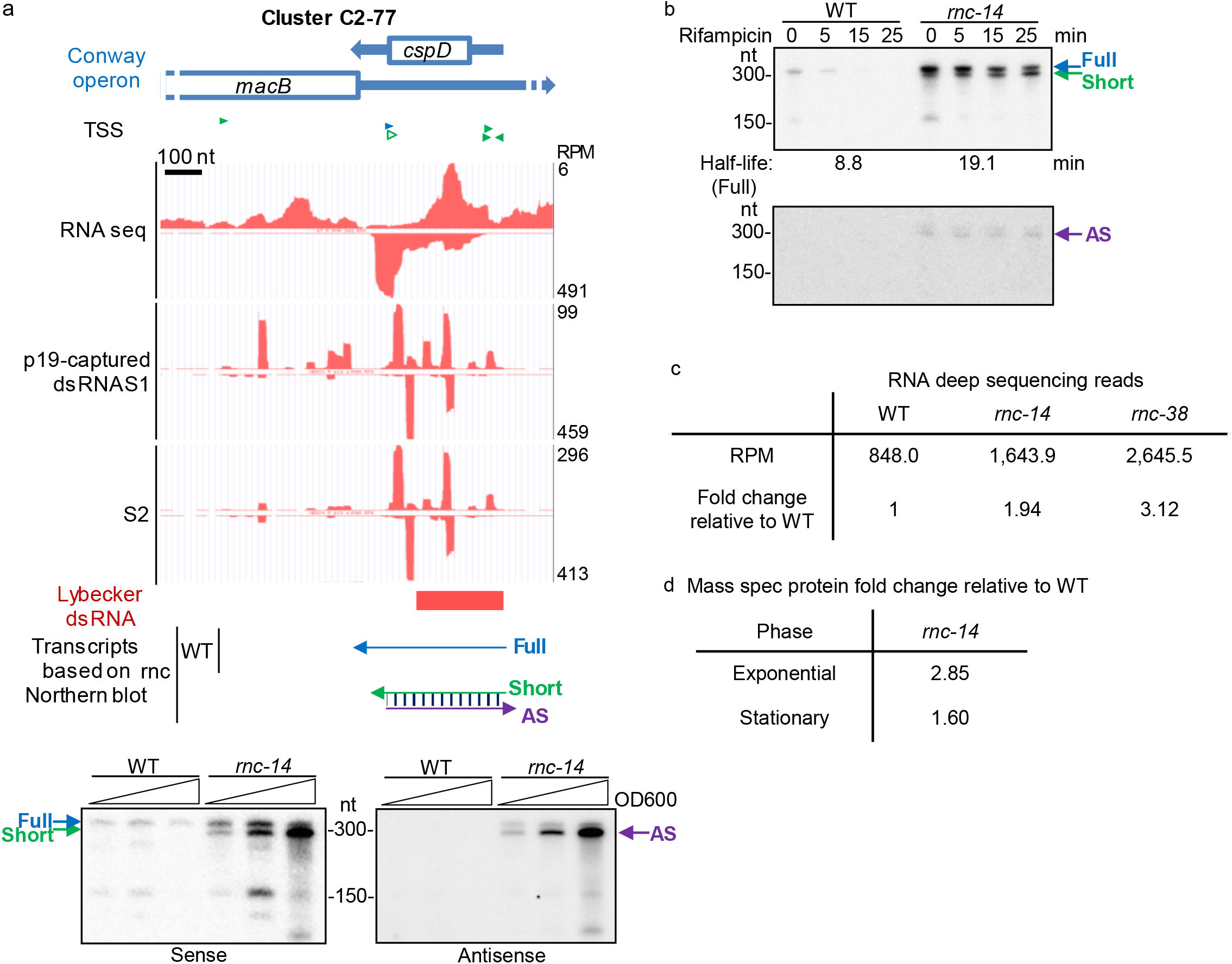
*cspD* is regulated by an antisense transcript and RNase III. a. Top: p19-captured dsRNA and total RNA sequencing data plotted in the UCSC genome browser; Middle: schematic of predicted RNA transcript architect at *cspD* locus; Bottom: Northern blots probing sense and antisense transcripts of *cspD.* Arrows indicate detected putative RNA transcripts with the color scheme matching that in the schematic. AS: antisense transcript. b. RNA half-life assay for sense and antisense transcripts of *cspD.* c. *cspD* mRNA abundance measured by RNA deep sequencing. d. CspD protein fold change (median of three replicates) measured by quantitative proteomics.

The last type of dsRNA involves dsRNAs arising from unannotated asRNA. Some of the p19-captured dsRNA loci could not be assigned to divergent gene transcripts or known asRNA, suggesting they arise from uncharacterized asRNA. One example is the C2-43 locus that contains both *yajO* and *dxs* genes on one strand (Fig. 4). RNA-seq reads, corroborated by an asTSS and Conway operon, suggest that there is an antisense transcript (opposite to *yajO* and *dxs)* that begins downstream of the 3’ end of *dxs*. p19-captured dsRNA reads in this cluster were adjacent to the beginning of the RNA-seq overlap, suggesting that RNase III cleavage might have helped to form the end of this asRNA. This asRNA could potentially pair with the 5’ UTR of *yajO* or 3’ UTR of *dxs*. Other examples include C2-169 in which the p19-captured dsRNA profile suggests an antisense transcript within the CDS of *galF* and C2-283, which predicts an antisense transcript in *secY* (Supplementary Fig. 7b).

### Confirmation and characterization of the *rsd* antisense transcript

To verify the sequencing results, the *rsd* gene, which was amongst the 3 most abundant coding gene p19-captured dsRNA clusters in both S1 and S2, was analyzed by Northern blotting (Supplementary Fig. 8). Antisense reads, which overlapped with the 5’-end of the *rsd* transcript by RNA-seq, may have originated from divergently oriented antisense transcripts that could be the transcript of an adjacent gene, *nudC*. dsRNA could have been formed between the 5’ UTR of a *nudC* transcript and the 5’-end of an *rsd* transcript. This locus resembles the “excludon” in *Listeria*, where two operons on opposite strands overlap at 5’ ends (25). A faint and smeary ~500 nt signal for *rsd* sense RNA (coding sequence is 477 nt) was detected in both WT and *rnc* mutant at similar levels (Supplementary Fig. 8a). Two more abundant shorter *rsd* sense transcripts and similarly sized antisense transcripts between 150 and 300 nt in length were detected only in the *rnc* mutant, suggesting that the sense and antisense RNAs formed dsRNAs that were degraded by RNase III. The *rsd* asRNA was less abundant in bacteria deficient in both RNase III and *rpoS*, which encodes a general stress response sigma factor that induces gene expression in stationary phase, suggesting that transcription of the asRNA may have been induced by RpoS.

Next, 5’ RACE was used to identify the TSS of the antisense transcript using *rnc* mutant bacteria. asRNAs with two potential asRNA TSS were cloned that were located opposite to the 5’ region of *rsd* (Supplementary Fig. 8b). A putative promoter sequence associated with the more upstream of the two potential asRNA TSS (corresponding to 3 of 4 clones) was identified and assessed in a *lacZ* reporter system, together with a construct with predicted inactivating mutations. As expected, the WT antisense promoter drove *lacZ* expression, and the promoter activity was reduced 2.5-fold in the promoter mutant (p-value<0.0001). To confirm our identification of the asRNA promoter, HA-tagged *rsd* reporter plasmids containing the WT or mutated antisense promoter (synonymous mutation for *rsd*) were introduced into WT and *rnc* mutant *E. coli* and expression of *rsd* sense and antisense transcripts and Rsd-HA protein were assessed by Northern blot and immunoblot, respectively (Supplementary Fig. 8c). Mutation of the antisense promoter reduced at least one of the short sense *rsd* transcripts and the two antisense transcripts. These findings confirm the identification of the antisense promoter of *rsd* asRNA. There was also a suggestion that the full-length Rsd transcript and protein levels might be slightly reduced. However, we cannot exclude the possibility that the antisense promoter mutation might affect the stability of the *rsd* sense transcript. These data suggest that Rsd might be subtly regulated by its asRNA, but we were unable to show that this difference has any biological significance on cell growth or survival.

### RNase III-dependent and asRNA regulation of CspD protein

Another coding gene with an abundant asRNA was *cspD*, a cold shock protein (CSP) family gene (which actually is not induced by cold shock in *E. coli*). CspD binds to DNA and can inhibit DNA replication (46). CSP proteins in *Salmonella* bind RNA and are involved in bacterial virulence (47). Although p19-captured dsRNA reads covered the entire CDS of *cspD* (225 nt), asRNA were detected by Northern blot only in the *rnc* mutant (Fig. 5a), suggesting that the *cspD* asRNA is not stable in WT cells. Both the level and half-life of the full length (~300 nt) *cspD* RNA increased in the *rnc* mutant, suggesting that *cspD* is a direct target of asRNA and that regulation depends on RNase III (Fig. 5a, b). A slightly shortened *cspD* sense RNA of the same size as the *cspD* asRNA was detected only in the *rnc* mutant. The length of the short sense RNA and asRNA were roughly equal to the length of the region covered by dsRNAs. These data suggest that dsRNAs containing the overlapping region of the sense and antisense RNAs accumulated in the *rnc* mutant. A *cspD* dsRNA was also identified by dsRNA antibody pulldown at approximately the same location (22). The RNA-seq reads of *cspD* RNA increased ~2-3-fold in *rnc-14* and *rnc-38* mutants (Fig. 5c and Supplementary Table 2), consistent with the Northern blots (Fig. 5a). Quantitative proteomics also found increased CspD in the *rnc-14* mutant (Fig. 5d and Supplementary Table 4). These data suggest that *cspD* mRNA and protein expression are reduced by asRNA in a RNase III-dependent manner.

### RNase III cleavage of overlapping sense-antisense RNAs does not have a consistent effect on protein abundance

To investigate if RNase III cleavage of overlapping antisense RNA affects protein levels, exponential and stationary phases of WT and *rnc-14 and rnc-38* cell lysates were analyzed by quantitative proteomics using the Tandem Mass Tag method (48). Approximately 400 proteins were identified with high confidence in both phases (Supplementary Fig. 9, Supplementary Table 4). When the relative changes in protein levels in each *rnc* mutant were compared to WT levels, changes in protein abundance in the mutants and the abundance of p19-captured dsRNAs in WT bacteria were not correlated. The levels of proteins encoded by genes producing abundant dsRNAs did not consistently increase in *rnc* mutants, although CspD increased (marked in red in Supplementary Fig. 9a). These findings suggest that RNase III cleavage of sense-asRNA duplexes does not globally impact protein abundance under homeostatic conditions.

To identify individual proteins that might be affected by RNase III deficiency, the ratio of protein abundance in both *rnc* mutant strains relative to WT in exponential phase was plotted (Fig. 6c, Supplementary Fig. 9b). Several proteins were consistently upregulated (YjhC, GabD AceA, AceB) or downregulated (SodA) in both *rnc* mutants in exponential phase samples. These proteins are involved in glycolysis and antioxidant responses.

**Figure 6.**
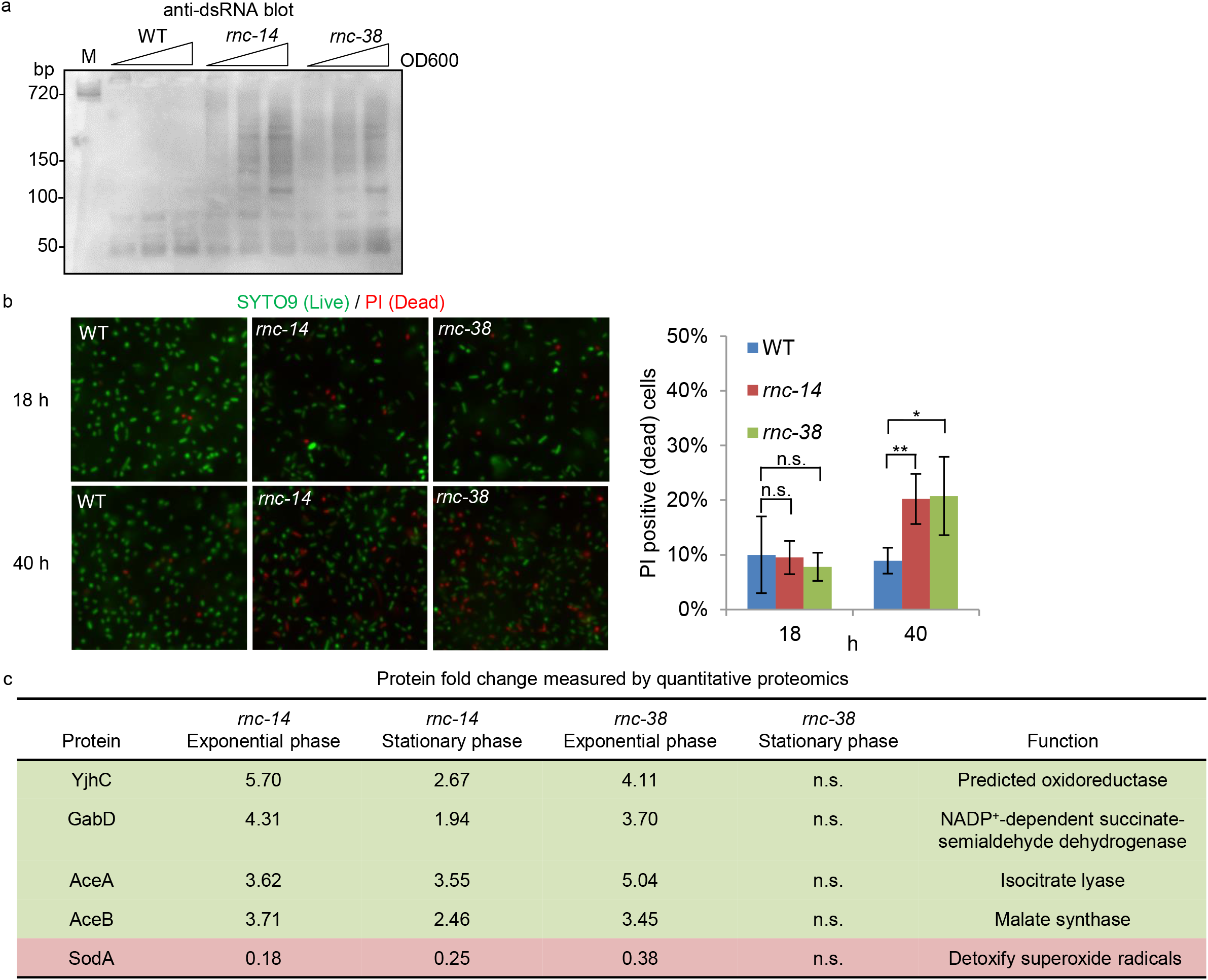
Accumulation of dsRNAs in RNase III mutant bacteria and physiological consequences. a. Anti-dsRNA immunoblot (probed with J2 anti-dsRNA) of total RNAs isolated from WT and *rnc* mutant *E. coli* during growth. Gel stained by SYBR-Gold before blotting is shown in Supplementary Fig. 1 b. M: dsRNA of 720 bp. b. Left: representative images of Live/Dead staining of WT and *rnc* mutant *E. coli* in two culture stages (18 or 40 h); right: quantification of the percentage of PI-positive dead cells. Data are from 4 biological repeats. Error bars show the standard deviation. n.s.: not significant; *: *p*-value<0.05; **: *p*-value<0.01. c. Selected proteins with consistent changes in both *rnc-14* and *rnc-38* mutants compared to WT (upregulated proteins in green and downregulated protein in red; also marked in Supplementary Fig. 9b). n.s.: not significant, which means protein fold change did not pass the cutoff for significance.

### RNase III mutants have increased cell death in late stationary phase

To begin to determine whether RNase III cleavage of dsRNA has biological significance, we used J2 antibody to assess the abundance of dsRNA during bacterial growth by immunoblotting equal amounts of electrophoresed RNA (Fig. 6a, Supplementary Fig. 1b). dsRNAs >100 bp in length accumulated in *rnc-14* and *rnc-38*, but not WT cells, as bacteria entered stationary phase. The relative amount of dsRNAs in the *rnc* mutant strains increased with cell density, suggesting that more dsRNAs were formed during stationary phase that were degraded by RNase III in WT cells. LIVE/DEAD staining of WT and *rnc* mutant strains showed no significant difference in death in early culture (18 hr) but about twice as much cell death in late stationary phase (40 hr, p-value<0.01 for *rnc-14* andp-value<0.05 for *rnc-38*, Fig. 6b). Thus, RNase III reduces dsRNA and protects cells from cell death in late stationary phase.

## Discussion

Here we developed a method to capture endogenous small dsRNAs (~21 bp) by ectopic expression of tombusvirus p19 in *E. coli*. Deep sequencing of p19-captured dsRNAs and total rRNA-depleted RNA suggested that clusters of short dsRNAs arise from long duplexes formed by overlapping sense and antisense transcripts that are processed into short dsRNAs by RNase III. p19 capture did not appear to introduce any substantial sequence bias, but stabilized labile dsRNA products to enable us to detect dsRNA with high sensitivity. asRNAs were transcribed from most genes, as previously noted (2, 22, 49), but with a wide-range of abundance. The abundance of captured dsRNAs correlated with asRNA reads. Although some of the less abundant asRNA and dsRNA may represent transcriptional noise, the most abundant p19-captured dsRNA clusters we identified agreed well with asRNA identified in other studies by deep sequencing, assignment of antisense transcription start sites (38) and operons (39), and dsRNA captured with anti-dsRNA antibody (22) and are likely the result of intended transcription. Our method confirmed hundreds of previously identified asRNAs and identified potentially hundreds of new such loci. One advantage of p19 capture is that it was performed in WT bacteria, potentially avoiding secondary effects caused by RNase III deficiency in RNase III mutant cells used in some studies (21, 22). Our data should provide a valuable resource for studying asRNA in *E. coli* and the method could be readily adapted to study asRNA in other species. The p19-captured dsRNA and RNA deep sequencing data have been formatted for convenient viewing in the UCSC genome browser (in Supplementary data file 1-4 in bedGraph format).

Many previous studies of the function of asRNA and RNase III in bacteria (2, 22, 49), including ours, utilized *rnc* mutant strains. RNase III degrades perfect dsRNAs generated from pairing of sense and antisense transcripts, but can also cleave structured RNAs that contain perfectly or imperfectly paired double-stranded regions (e.g. rRNA precursor (31) and R1.1 RNA of T7 phage (50)). Therefore, some changes in RNA level and half-life in *rnc* mutant bacteria are due to RNase III degradation of dsRNAs generated from pairing of sense and antisense transcripts, but others may be due to RNase III cleavage of structured RNAs. There is no simple way to separate the antisense dependent effects of RNase III. However, p19 only binds perfectly paired dsRNAs, such as would arise from antisense transcripts pairing with sense transcripts, but not imperfect duplexes that would arise in structured regions of RNA that might also be substrates of RNase III, providing a specific way to capture antisense transcripts. Supplementary Table 5 shows a comparison of our method with previous methods that have identified RNase III targets (2, 21, 22, 49, 51, 52). Our method is the only method that can be performed in WT cells and reveal exact RNase III cleavage sites.

The most abundant p19-captured dsRNA clusters, which were mostly found in other studies and least likely to be caused by transcriptional noise, were confirmed by Northern blotting and studied in more detail. Most of these clusters arose from divergent transcripts from overlapping 5’ regions of operons on opposite strands. RNase III deficiency for most of the abundant clusters increased the quantity and/or half-life not only of the antisense transcript, but also of the corresponding sense transcript, suggesting that asRNA transcription and RNase III degradation of dsRNAs promote more efficient sense RNA decay. Moreover, shorter asRNAs were generally detected only in RNase III-deficient bacteria. dsRNAs accumulated as RNase III-deficient bacteria exited exponential growth in stationary phase when RNase III-deficient bacteria were significantly more prone to undergo cell death. However, despite widespread antisense transcription, unbiased quantitative proteomics did not indicate global changes in protein levels that correlated with the abundance of dsRNAs at a locus. This finding suggested that antisense transcription might not play a large role in regulating protein levels and bacterial physiology under most conditions. However, a few proteins associated with antioxidant responses and glycolysis reproducibly were altered in two RNase III mutant strains and the mRNA and protein level of CSP CspD increased in the mutant strains. These proteins might be more important in stressed conditions. Moreover, many of the most abundant p19-captured dsRNA clusters, including *ryeA/ryeB*, the toxin-antitoxin genes, and tRNAs, might also be particularly important during stress. These findings, when considered with the increased death in late stationary phase of RNase III mutant bacteria, suggest that asRNAs and degradation of dsRNAs, especially those regulating noncoding RNAs, might only become important when nutrients are limiting or during other forms of environmental stress. However, we cannot exclude the possibility that RNase III activity on structured RNAs could be responsible for the increased death of *rnc* mutants in late stationary phase. Additional work is required to determine whether antisense transcription has any physiological role and under what conditions.

A few well studied small regulatory RNAs generate abundant dsRNAs (Table 1). Our data demonstrate that RNase III cleaves all three families of cis-acting Type I TA class small RNAs including *ldrD/rdlD, mokC/sokC*, and *ibsD/sibD* (Fig. 3d, Supplementary Fig. 6), suggesting that one way RNase III may regulate bacterial survival and physiology is by controlling toxic proteins. The abundance and stability of the *spf* small RNA were also shown to be regulated by RNase III. Its maturation might require RNase III processing (Fig. 3c). Small RNAs, like *spf* and *micA*, function *in trans* to regulate many other genes and their functions could require the RNA chaperone Hfq (53). RNase III may indirectly impact gene expression by regulating important small regulatory RNAs like *spf*.

Although asRNA and/or RNase III cleavage may not be important for regulating sense gene translation into protein under homeostatic conditions, RNase III could still be important for eliminating paired sense and antisense RNAs and structured RNA fragments. For most coding and non-coding genes we examined, RNA half-lives were increased in *rnc* mutants (Fig. 3, 5). For many of the abundant p19-captured dsRNA clusters, shortened RNA fragments and dsRNAs were detected only in *rnc* mutants (Fig. 3, 5, Supplementary Fig. 6, 8). These findings might best be explained if RNase III cleavage acts downstream of other RNases, which produce shortened RNAs as decay intermediates (model in Supplementary Fig. 10). RNase E is a 5’-end-dependent (54) single-stranded endoribonuclease that acts on most *E. coli* mRNAs. After the mRNA is cut, the newly formed 3’-end can then be degraded by PNPase, which also acts only on single-stranded and nonstructured RNAs. PNPase would stall at 5’ ends of overlapping sense and antisense transcripts or at structured regions. We propose that RNase III clears these stalled dsRNAs. The shortened RNA products that accumulate in *rnc* mutants in small RNA loci like *ryeA/ryeB* (Fig. 3b) or in coding genes like *rsd* (Supplementary Fig. 8a), might be stalled products of RNase E and PNPase that are further degraded by RNase III. RNase III prefers to cleave dsRNAs into ~14-bp or shorter fragments, which can then be completely degraded into mononucleotides by the concerted actions of RNA helicases, PNPase and oligoribonuclease (55, 56). Lack of clearance of dsRNA intermediates, which accumulate in stationary phase in *rnc* mutants, could be toxic and lead to increased death of *rnc* mutant bacteria in this phase. dsRNAs are also sensed by innate immune receptors in eukaryotes, which might be costly for bacteria during infection. However, further work is required to explore these conjectures.

*cspD* appears to be a rare example of RNase III-regulated protein production through a *cis-* acting asRNA (Fig. 5). RNase III might be essential for degrading *cspD* sense mRNA. *cspD* asRNA covers a substantial region of the sense RNA and the dsRNA might mask cleavage sites of other RNases (e.g. RNase E) and stabilize the *cspD* sense RNA. A similar mechanism in which asRNA stabilizes sense RNA and impedes RNase degradation has been described for the *gadY* small RNA, which stabilizes overlapping *gadX* mRNA (57).

p19 capture could be used to identify RNase III cleavage sites to better understand RNase III mechanism and sequence preferences. Bacterial RNase III is thought to recognize structural features (A-form helix) of dsRNA rather than a sequence motif (58). However studies on RNase III targets in cells clearly show selection for certain genes (49, 55), suggesting RNase III might have sequence bias. So far only weak consensus sequences for RNase III recognition were identified based on RNase III cleavage of a small number of structural RNAs (55, 59). At least one G/C pair is preferred for RNase III cleavage of substrates with stem loop structure in one study (49). However, the dsRNA hot spot patterns were identified *in vitro* and in cells (Supplementary Fig. 2, (28)), strongly suggest that *E. coli* RNase III has some intrinsic sequence bias. We previously demonstrated that p19-captured dsRNA hot spot patterns were not due to cloning bias (28). Surprisingly, cleavage bias also seemed likely for human Dicer in our *in vitro* digestion assay (E4 in Supplementary Fig. 2), although current models suggest that Dicer cuts dsRNAs from the 3’-end and in a phased manner without bias (60). However, a few recent studies have shown sequence preferences for RNase III class enzymes, including Mini-III in *Bacillus subtilis* (61), yeast Rnt1p (62), and dicer-like enzymes in *Paramecium* (63). A GC bias was also found in plant viral-derived siRNAs (64). Further analysis of dsRNAs in *E. coli* and other bacterial species may help to unravel the mechanisms underlying sequence bias of RNase III class enzymes. Of note, RNase III is required for making guide RNAs for the bacterial CRISPR system and any sequence bias of RNase III could potentially influence the selection of genes targeted by CRISPR.

In summary, our study presents a new method for identifying and studying asRNA in bacteria that could also be adapted to eukaryotic studies. p19-captured dsRNA clusters mark genomic loci where overlapping sense and antisense transcription occur in *E. coli*. p19-captured dsRNAs are most prominent at the 5’-end of genes, where divergent transcription of sense and antisense transcripts is widespread. Similar divergent antisense transcripts overlapping at the 5’-end of mRNAs have also been described in eukaryotes (65). Failure to clear dsRNAs, which requires RNase III, may be toxic to cells, especially during environmental stress. Although abundant asRNAs regulate sense RNA decay, their effect on protein expression appears to be subtle. More work is needed to understand the role of asRNA in bacteria and the consequences of not efficiently clearing the dsRNAs that form.

## Materials and Methods

Bacterial strains, plasmids, and oligonucleotides used in this study are listed in *SI Materials and Methods* and Supplementary Table 6. Ectopic expression of p19 and dsRNA isolation were based on our previous methods (28, 29) with modifications described in *SI Materials and Methods*. Detailed protocols for bacterial culture, total RNA extraction, small RNA and total RNA deep sequencing, bioinformatic analysis, RNA immunoblot, Northern blot, Western blot, 5’ RACE, *lacZ* reporter assay, proteomics, and statistics are included in *SI Materials and Methods*.

## Supporting information

Supplementary Table 1. Abundance of p19-captured dsRNAs in E. coli genes

Supplementary Table 2. Total RNA sequencing data for WT and rnc mutant E. coli

Supplementary Table 3. p19-captured dsRNA clusters

Supplementary Table 4. Quantitative proteomics comparing WT and rnc mutants

Supplementary data file 1. p19-captured dsRNA S1 sense strand reads in bedGraph format

Supplementary data file 2. p19-captured dsRNA S1 antisense strand reads in bedGraph format

Supplementary data file 3. p19-captured dsRNA S2 sense strand reads in bedGraph format

Supplementary data file 4. p19-captured dsRNA S2 antisense strand reads in bedGraph format

## Acknowledgements

We thank Sidney Kushner for providing MG1693 and SK7622 strains. L.H. is supported by a NSFC grant (31870128), a Shenzhen Science and Technology grant (JI20170301), and a GSK-IDI Alliance Fellowship.

## Conflict of interest

J.J. and L.M. are employees of New England BioLabs (NEB), which sells p19 protein, enzymes, sequencing library construction kits, and other research reagents. Other authors declare no conflict of interest.

## Supplementary Materials and Methods

### Bacterial strains, plasmids, and culture conditions

*E. coli* strain MG1693 and its derivative, SK7622 *(rnc-38* mutant), were utilized in the experiments with pcDNA3.1-p19-FLAG plasmid (Supplementary Table 6). MG1655 and MG1655 *ΔlacZYA* (also referred to as MG1655 *Δlac)*, and derivatives with mutations in *rnc*, or *rnc* and *rpoS*, and the chromosomal *p19* expression construct, are described in Supplementary Table 6. *E. coli* strain FW102 was used to construct the single copy *rsd* antisense promoter-lacZ fusions (Supplementary Table 6). Plasmids for expressing p19-FLAG and Rsd-HA and for synthesizing dsRNAs are described in Supplementary Table 6. Unless indicated, strains were cultured in LB (Lennox, BD) at 37°C with shaking at 250 rpm and antibiotics when required were used at the following concentrations: carbenicillin (100 μg/ml), kanamycin (10 or 25 μg/ml), and tetracycline (12.5 μg/ml).

### *E. coli* total RNA extraction

For each 5 ml of *E. coli* culture, 5 ml of cold methanol was added to the sample immediately after harvesting in order to stabilize RNA, and the sample was kept on ice for processing. After centrifugation, the bacterial pellet was resuspended in 1 ml lysis buffer (4 M guanidinium thiocyanate, 25 mM sodium citrate, pH 7.0, 0.5% (wt/vol) N-laurosylsarcosine (Sarkosyl) and 0.1 M 2-mercaptoethanol) (1). To ensure complete disruption of bacterial cells, samples were processed in a bead beater (Biospec) with glass beads. The lysate was centrifuged at 20,000 g for 30 min and RNA was extracted from the cleared lysate using the protocol of Chomczynski and Sacchi (1). DNA contamination was removed by DNase I digestion (M0303L, NEB) and RNA was purified using acid-Phenol:Chloroform (AM9722, Invitrogen) according to the manufacturer’s protocol.

### Extraction of p19-captured small dsRNAs in *E. coli*

p19 capture of dsRNAs was performed on WT *E. coli* (MG1693) and *rnc-38* (SK7622) transformed with pcDNA3.1-p19-FLAG after overnight culture. For the *E. coli* strain with the genome-integrated *p19* gene, an overnight culture of *E. coli* was diluted 200 times to inoculate fresh broth. In the case of the S1 exponential phase sample, when the culture reached an OD_600_ of 0 4, isopropyl-β-D-thiogalactoside (IPTG) was added at 0.5 mM for 1 h (final OD_600_ of the culture was 1.2). In the case of the S2 stationary phase sample, when the culture reached an OD_600_ of 1.4, IPTG was added at 0.5 mM for 1 h (final OD_600_ of the culture was 2.0). Total RNAs were extracted as described above and p19 magnetic beads (from p19 miRNA Detection Kit, E3312, NEB) were used to pull down small RNAs from total RNAs (isolated from 20 ml of bacterial culture) as previously described (2).

### Northern blotting

Northern blotting was performed using two methods. Method 1 used a 5% TBE-Urea polyacrylamide RNA gel cast using the Bio-Rad Mini-PROTEAN Tetra Cell system (2). RNA samples (15 μg total RNA) were heated to 95°C for 5 min in Gel Loading Buffer II (AM8546G, Ambion) and immediately placed on ice until gel loading. Electrophoresis was performed at room temperature and the gel was run at 150 V for about 1 h. Gels were stained with SYBR-Gold (S11494, Invitrogen) and then transferred to a Hybond-N+ Membrane (RPN303B, Amersham) by capillary transfer in 20X SSC buffer (AM9763, Ambion) overnight. The membranes were UV crosslinked. Low Range ssRNA Ladder (N0364S, NEB) was used as size markers. Blots for *ryeA/ryeB, spf, ldrD/rdlD, cspD, micA, mokC/sokC*, and *ibsD/sibD* loci transcripts were performed using Method 1.

Method 2 used denaturing formaldehyde agarose gels (1.2%) in MOPS buffer (AM8671, Ambion) electrophoresed in a mid-sized horizontal gel tank at 80 V for about 4 h at room temperature. Other procedures were as in Method 1. The ssRNA Ladder (N0362S) from NEB was used as the size standard. Blots for the *rsd* locus transcripts were performed using Method 2.

DNA oligos were obtained from IDT. The DNA oligo probes were: for *ryeA/ryeB* locus, probe for *ryeA:* 5’-CAACTTTTAGCGCACGGCTCTC-3’, probe for *ryeB:* 5’-GAGACCGAACACGATTCCTGTA-3’; for *spf* locus, probe for sense transcript: 5’-TAAAAAACGCCCCAGTCATTACTGACTGGGGCGGCTAAAATATTCAGCCA-3’, probe for antisense transcript: 5’-TGGCTGAATATTTTAGCCGCCCCAGTCAGTAATGACTGGGGCGTTTTTTA-3’; for *ldrD/ rdlD* locus, probe for *ldrD:* 5’-AGTGGTCTAGAGTCAAGATTAGCCCCCGTGGTGTTGTCAGGTGCAT-3’, probe for *rdlD*: 5’-AGAAAACCCCCGCACGTTGCAGGTATGCACCTGACAACACCACGGG-3’; for *cspD* locus, probe for sense transcript: 5’-GAACGGATTGTCCAGCTTTTAGCGTTCTGT-3’, probe for antisense transcript: 5’-ACAGAACGCTAAAAGCTGGACAATCCGTTC-3’; for *micA* locus, probe for sense transcript: 5’-AGGCGAGTCTGAGTATATGAAAGACGCGCATTTGTTATCATCATCCCTGA-3’, probe for antisense transcript: 5’-TCAGGGATGATGATAACAAATGCGCGTCTTTCATATACTCAGACTCGCCT-3’; for *mokC/sokC* locus, probe for *mokC:* 5’-CCCTCTGATTGGCTGTTAATAAGCTGCGAAACTTACGAGTAACAACACA-3’, probe for *sibD*: 5’ -TGTGTTGTTACTCGTAAGTTTCGCAGCTTATTAACAGCCAATCAGAGGG-3’; for *rsd* locus, probe 1 for sense transcript (used in Supplementary Figure 8a): 5’-AGTTTGTTACTTCCTCTGACGCGCTCCGTCAGGTTATCGAGCTGG-3’, probe 2 for sense CCAGCTCGATAACCTGACGGAGCGCGTCAGAGGAAGTAACAAACT-3’.

The DNA probes were 5’ end-labeled with γ-^32^P ATP (PerkinElmer) and T4 Polynucleotide Kinase (M0201L, NEB). For probe hybridization, the membrane was incubated with rotation in a hybridization oven in hybridization buffer (ULTRAhyb-Oligo, Ambion) at 42°C overnight. The membrane was washed 3 times, for 20 min each time, in 0.5% SDS, 2X SSC buffer (AM9763, Ambion) at 42°C with rotation in a hybridization oven. The membrane was visualized using a phosphorimager screen and FLA-9000 Image Scanner (Fujifilm).

For re-blotting a membrane with a second probe, the membrane was rotated in a hybridization oven in stripping buffer (0.1% SDS) at 90°C for 30 min. The probe stripping was verified by visualizing the membrane, and then the membrane with processed with the second probe.

### RNA half-life assay

WT and *rnc* mutant *E. coli* were cultured at 37°C with shaking at 250 rpm overnight and then diluted 200 times to start a fresh culture. After the cultures reached OD_600_ of ~0.5, a sample was extracted (termed time 0). Then, rifampicin in DMSO was added to a final concentration of 500 μg/ml and the culture growth was continued. Additional samples were harvested from the culture at 2, 5, and 12.5 min, or 5, 15, and 25 min. Upon harvest of each sample, one volume of cold methanol was immediately added to stabilize the RNAs. The samples were kept on ice and total RNA extraction and Northern blotting were performed as described above. Multi-gauge software (Fujifilm) and Image Studio Lite software (LI-COR) were used to quantify hybridization signals. RNA half-lives were calculated using the slope of a linear trendline fitted from the normalized intensity of hybridization bands.

### *E. coli* total RNA deep sequencing

Total RNAs were extracted as described above and ribosomal RNAs were removed using bacterial Ribo-Zero rRNA Removal Kit (MRZMB126, Epicentre) following the manufacturer’s protocol. RNA sequencing libraries, created using NEBNext Ultra Directional RNA Library Prep Kit for Illumina (E7420S, NEB) according to the manufacturer’s protocol, were sequenced on an Illumina GAII sequencer at NEB.

### *E. coli* RNase III and human Dicer *in vitro* digestion assay

To produce dsRNAs as the substrate for RNase digestion, the entire *eGFP* coding sequence (720 bp, from pEGFP-N1, Clontech), or a 523 bp fragment (nt 267 to 789) of the *LMNA* coding sequence (NM_005572.3), were cloned with the T7 promoter sequence flanking the 5’-ends of the DNAs. Sense and antisense RNAs were transcribed separately using T7 RNA polymerase (M0251L, NEB) according to the manufacturer’s protocol. Purified sense and antisense RNAs were mixed and annealed by heating to 90°C for 2 min and then gradually cooled at room temperature. Annealed RNA products were separated on a 6% native PAGE gel and isolated after staining with SYBR-Gold. dsRNAs were eluted overnight from gel pieces in 0.3 M NaCl. RNAs were precipitated by ethanol and then dissolved in nuclease-free water.

For each digestion reaction, ~200 ng of PAGE-purified dsRNAs were used and the resulting digestion products were analyzed by deep sequencing. For RNase III digestion, dsRNAs were incubated for 20 min at 37°C with ShortCut RNase III (M0245L, NEB) in 1X digestion buffer supplemented with either 10 mM MgCl_2_ or 20 mM MnCl_2_. p19 magnetic beads (NEB) were used to pulldown small dsRNA products of the RNase III digestion reaction buffer supplemented with MnCl_2_. Recombinant (human) Dicer Enzyme Kit for RNA Interference (Genlantis) was used for Dicer digestion. The digestion reaction was carried out at 37°C overnight (for ~20 hours), performed according to the manufacturer’s protocol.

### Small RNA cloning and deep sequencing

p19-captured small dsRNAs isolated from *E. coli* cells expressing p19 from a plasmid were cloned and sequenced according to Huang et al. (2). p19-captured small dsRNAs isolated from *E. coli* cells with integrated *p19* were cloned using the NEBNext Small RNA Library Prep Set for SOLiD (E6160, NEB) according to the manufacturer’s protocol and the small RNA libraries were sequenced on a SOLiD sequencer at NEB. Small RNAs from RNase III and Dicer digestion assays were cloned using the NEBNext Small RNA Library Prep Set for Illumina (E7330L, NEB) according to manufacturer’s protocol and libraries were sequenced on an Illumina GAII sequencer at NEB. All small RNA deep sequencing data are available under BioProject PRJNA512059 at Sequence Read Archive database, NCBI (http://www.ncbi.nlm.nih.gov/).

### Bioinformatic analysis

Cutadapt (https://cutadapt.readthedocs.io/en/stable/index.html) was used to trim cloning adapter sequences. Novocraft (www.novocraft.com) was used for sequence alignment using *E. coli* K12 MG1655 genome sequence (GenBank accession: U00096.2) and pcDNA3.1+ plasmid sequence (GenBank accession: EF550208.1) as references. A summary of sequence alignment results is included in Supplementary Table 7. SAMtools (http://samtools.sourceforge.net) was used to calculate sequencing reads for each gene and for generating sequencing profiles for both plasmid and genome. The UCSC genome browser *(E. coli* K12 Assembly: eschColi_K12, http://microbes.ucsc.edu) was used to view sequencing data and other published datasets. The p19-captured dsRNA S1 and S2 datasets were formatted into bedGraph files (Supplementary data file 1-4), which can be viewed directly using UCSC genome browser. Customized Perl scripts were created for small dsRNA cluster analysis and for formatting the datasets. All Perl scripts are available upon request.

### Western blot

Protein samples were prepared by resuspending bacterial cells in 1X SDS loading buffer and heating at 95°C for 5 min before SDS-PAGE. Antibodies and their dilutions were: anti-HA 1:1,000 (Sigma-Aldrich) and anti-His tag 1:500 (Covance, MMS-156P). Horseradish peroxidase conjugated anti-mouse or ant-rabbit IgG secondary antibodies were used at 1:10,000 dilution. SuperSignal West Pico Chemiluminescent Substrate (34580, Thermo Scientific) was used for developing ECL signals.

### RNA immunoblot

Total RNA (10 μg) was separated by native electrophoresis using mini-sized homemade 5% polyacrylamide TBE gels and a Bio-Rad Mini-PROTEAN Tetra Cell system. RNA samples were prepared in Gel Loading Buffer II (AM8546G, Ambion) and electrophoresed at room temperature. RNA was blotted onto a Hybond-N+ Membrane (RPN303B, Amersham) by capillary transfer overnight, and then UV-crosslinked. The membrane was first incubated with anti-dsRNA J2 antibody (used at 1:1,000, Scicons) in PBS buffer containing 5% BSA overnight at 4°C. HRP-conjugated anti-mouse secondary antibody was used at 1:10,000 and the signal was visualized using SuperSignal West Pico Chemiluminescent Substrate (34580, Thermo Scientific).

### 5’ rapid amplification of cDNA ends (RACE)

Total RNA isolated from the *E. coli* MG1655 *Δlac rnc-38* mutant was used for 5’ RACE to identify the antisense transcript at the *rsd* locus. Total RNA (2 μg) was treated with Antarctic phosphatase (M0289L, NEB) to remove the 5’ triphosphate and then with T4 polynucleotide kinase (M0201L, NEB) to add a 5’ monophosphate. A synthetic 5’ RNA adapter (5’-GUUCAGAGUUCUACAGUCCGACGAUC-3’) was ligated to RNAs containing the 5’ monophosphate using T4 RNA ligase (M0204L, NEB). cDNA was synthesized by SuperScript III reverse transcriptase (18080093, Invitrogen) using Rsd-F primer (5’-AGCGCGTCAGAGGAAGTAAC-3’). Two rounds of PCR were performed on the cDNA to amplify DNA fragments containing potential ends of *rsd* antisense transcript. Forward primer was from the sequence of the 5’ RNA adapter (5’-GTTCAGAGTTCTACAGTCCGA-3’). The reverse primer was Rsd-F2 (5’-GGAGCGCGTCAGAGGAAGTAAC-3’). A nested PCR reaction was performed using the products from the first round of PCR as the template and Rsd-F4 (5’-GGTTGATCGCTGGCTACATGTAC-3’) as the reverse primer. The product of the nested PCR was cloned into the pGEM-T easy vector (Promega) and multiple clones were sequenced by Sanger sequencing.

### β-galactosidase promoter reporter assay

FW102 *pASrsd-lacZ* and FW102 *pASrsd-mut-lacZ* reporter strain cells (Supplementary Table 6) were cultured in quintuplicate in 200 μl of LB in a 96 well plate at 900 rpm, 80% humidity, 37°C, in a multitron shaker. At an OD_600_ of 0.6, cells were lysed and processed for the β-galactosidase assays using microtiter plates and a microtiter plate reader, and Miller units were calculated as described (3).

### Protein quantification

#### Preparation of Protein Extracts

For each growth phase, three TMT (Tandem mass tag) 6-plex experiments were run consisting of WT, *rnc-14, rnc-38, hfq* bacteria and a pool of equal amounts of all of the samples in the same phase (20 ug protein of each sample) and a pool of the samples in the stationary and exponential phase (10 ug protein of each sample). The pool of samples across the phases used to normalize to their respective samples was not utilized in this experiment. Cells were lysed in PBS with HALT protease inhibitor cocktail (78430, Thermo Scientific) using a bead beater with glass beads. The protein concentration was determined using Pierce BCA protein assay reagent (23227, Pierce). Each strain was prepared in biological triplicate; 80 μg of protein from each sample was reduced with 20 mM dithiothreitol (DTT, Thermo Scientific) at 37°C for 1 hour and then alkylated using 50 mM iodoacetamide (IAM, Sigma) in 50 mM triethylammonium bicarbonate (TEAB, Sigma) at room temperature for 1 hour in the dark. The samples were then digested overnight at 37°C with sequencing grade (1:50) trypsin (V5111, Promega, Madison, WI). The samples were acidified with formic acid (FA) and lyophilized using a SpeedVac. The samples were re-suspended in 30 μl 500 mM TEAB. TMT (90063, Thermo Scientific, San Jose, CA), resuspended at room temperature in 70 μL acetonitrile (ACN), was added to each sample at room temperature for 1 hour and the reaction was stopped with 10% hydroxylamine. Samples from all 6 channels were combined and cleaned up using the Oasis HLB elution plate 30 μM (186001828BA, Waters, Milford, MA). The samples were lyophilized and reconstituted in 20 μL of 20 mM ammonium formate (pH 10), 2% ACN. All samples were then fractionated using a previously described high pH fraction method (4) into 12 fractions using a Dionex HPLC (Thermo Scientific, San Jose, CA) and 2.1 x 50 mm Xterra column (186000408, Waters, Milford, MA). The fractions were lyophilized and stored at −20°C until mass spectrometry analysis.

#### Mass Spectrometry

The tryptic peptides were reconstituted in 10 μL 2% ACN, 0.2% FA. The sample was first loaded at 5 μL/min onto a u-Precolumn 300 μm i.d. x 5 mm C18 PepMap100, 5 μm, 100Å trap column (160454, Thermo Scientific). Digested samples (2 μL) were analyzed using nanoflow liquid chromatography coupled to a data dependent mass spectrometer (LC/MS-MS) using the Eksigent nano-LC (Applied Biosystems/MDS Sciex, Foster City, CA) coupled to an LTQ-Orbitrap-Velo mass spectrometer (Thermo Scientific, San Jose, CA). A 75 μm id Picotip emitter with a 15 μm diameter tip (PF360-75-15-N-5, New Objective, Woburn, MA) was hand packed using Magic C18 100Å 3 μm resin to a length of 13 cm. Tryptic peptides were eluted over a 68 min gradient at a flow rate of 400 nL/min using a water/ACN gradient (Mobile Phase A: 100% water, 0.2% FA; Mobile Phase B: 100% ACN, 0.2% FA). The gradient was ramped from min 2% B to 40% B over 68 minutes, then ramped to 95% B over 8 min, held for 2 min at 95% B, ramped to 5% B in 2 min and then ramped to 2% B for 5 min.

The Velos system was operated in the standard scan mode with positive ionization. The electrospray voltage was 2.75 kV and the ion transfer tube temperature was 300°C. Full MS spectra were acquired in the Orbitrap mass analyzer over the 350-2000 m/z range with mass resolution at 60,000 (at 400 m/z); the target value was 2.00E+05. The 10 most intense peaks with a charge state greater than or equal to 2 were fragmented in the HCD collision cell with normalized collision energy of 40%. The tandem mass spectra were acquired in the Orbitrap mass analyzer with mass resolution of 120,000 with a target value of 1.00E+05. Ion selection threshold was 500 counts and the maximum allowed ion accumulation time was 100 ms for full scans. Dynamic exclusion was enabled with a repeat count of 1, repeat duration of 15 s, exclusion list of 500 and exclusion duration of 15 s. All samples were analyzed in biological triplicate and subjected to duplicate LC-MS/MS analysis.

#### Data processing and Protein Identification

Protein sequences from *E. coli* (strain K12) were obtained from www.uniprot.org on May 21 2013, appended with its own reversed sequences and with common mass spectrometry contaminant protein sequences and used for peptide and protein identification (9658 sequences; 3093176 residues). Raw data from the LTQ-Orbitrap-Velos were processed with Mascot (vs 2.2) using default parameters. The data were searched using trypsin as the enzyme and allowing for up to 2 missed cleavages. The search criteria included peptide mass tolerance (± 15 ppm), fragment mass tolerance (± 0.05 Da), fixed modifications of Carbamidomethyl (C) and variable modifications: Oxidation (M), Phospho (ST), Phospho (Y), TMT6plex (K) and TMT6plex (N-term). Mass values were monoisotopic and protein mass was unrestricted. Mascot results for sample fractions were aggregated and submitted to the PeptideProphet and ProteinProphet algorithms for peptide and protein validation, respectively (ISB/SPC Trans Proteomic Pipeline TPP v4.3 JETSTREAM rev 1, Build 200909091257 (MinGW)). Protein results were then filtered using a false discovery rate of <1%.

#### Protein Quantification and Statistical Analysis

The TMT reporter channel ion intensities were summed by peptide sequence with isotopic correction factors applied per manufacturer’s guidelines. Peptide fold changes were calculated across WT and mutant strains in exponential and stationary phase. The peptide fold changes were normalized using the median fold change of all quantified peptides. The protein fold-changes were derived from median peptide fold changes. Significance was determined using ANOVA statistical testing and *p*-values were calculated. The data was sorted using an adjusted *p*-value of 0.2 in a minimum of two of the three biological samples compared to the respective wildtype. Median protein fold-changes of the three biological replicates were calculated (Supplementary Table 4). The full data set has been deposited to the Proteome Xchange Consortium (http://proteomecentral.proteomexhange.org) via the PRIDE partner repository (5), with the identifier PXD011180.

### Live/Dead staining

The LIVE/DEAD BacLight Bacterial Viability Kit (L7007, Invitrogen) was used for staining WT and *rnc* mutant *E. coli* cells according to manufacturer’s protocol. Samples were harvested at 18 and 40 h after starting a fresh culture in LB medium.

### Statistics

Significance of differences between two samples was calculated using Student’s T-test. Significance of the difference between two correlation coefficients, based on Fisher r-to-z transformation, was calculated using an online tool: http://vassarstats.net/rdiff.html.

---

**Supplementary Figure 1.**
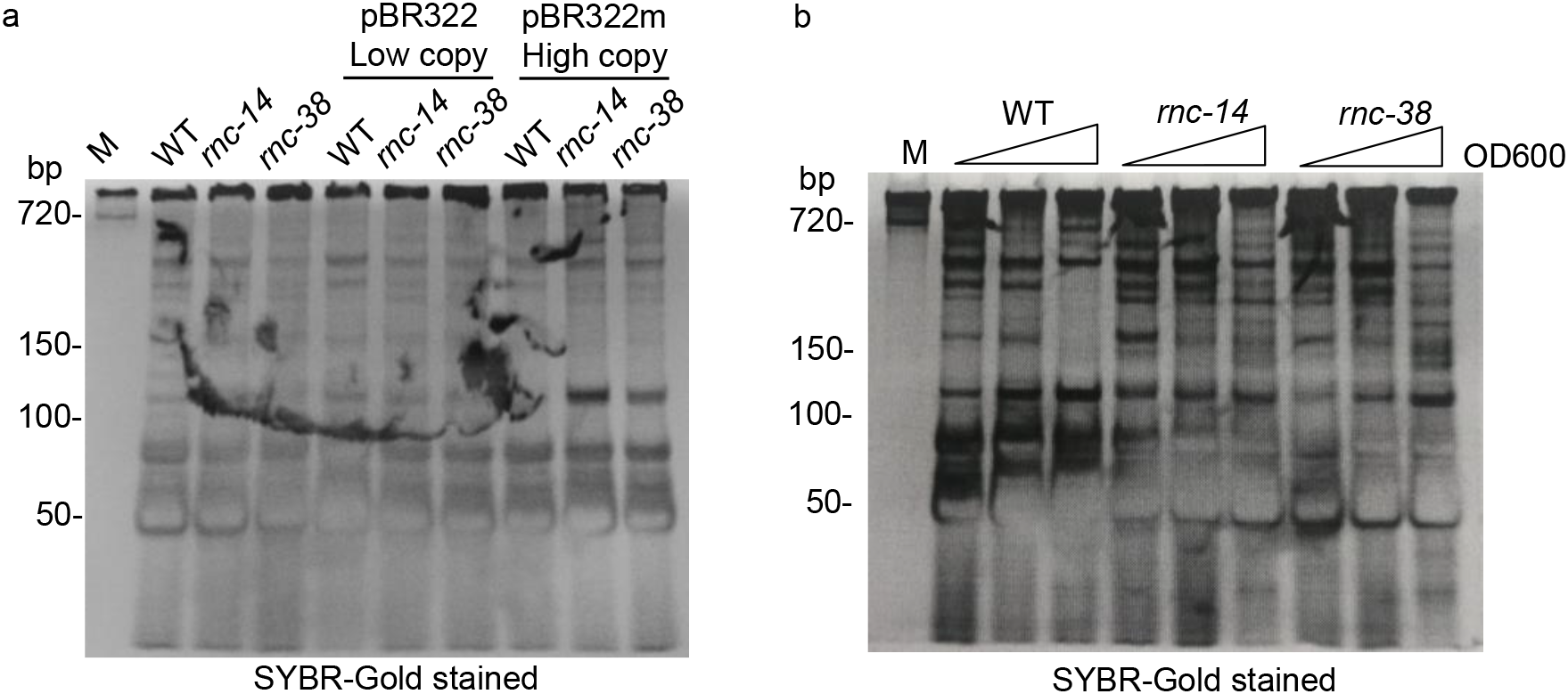
SYBR-Gold stained gels for anti-dsRNA blots. a. Same gel as Fig. 1e. b. Same gel as Fig. 6a.

**Supplementary Figure 2.**
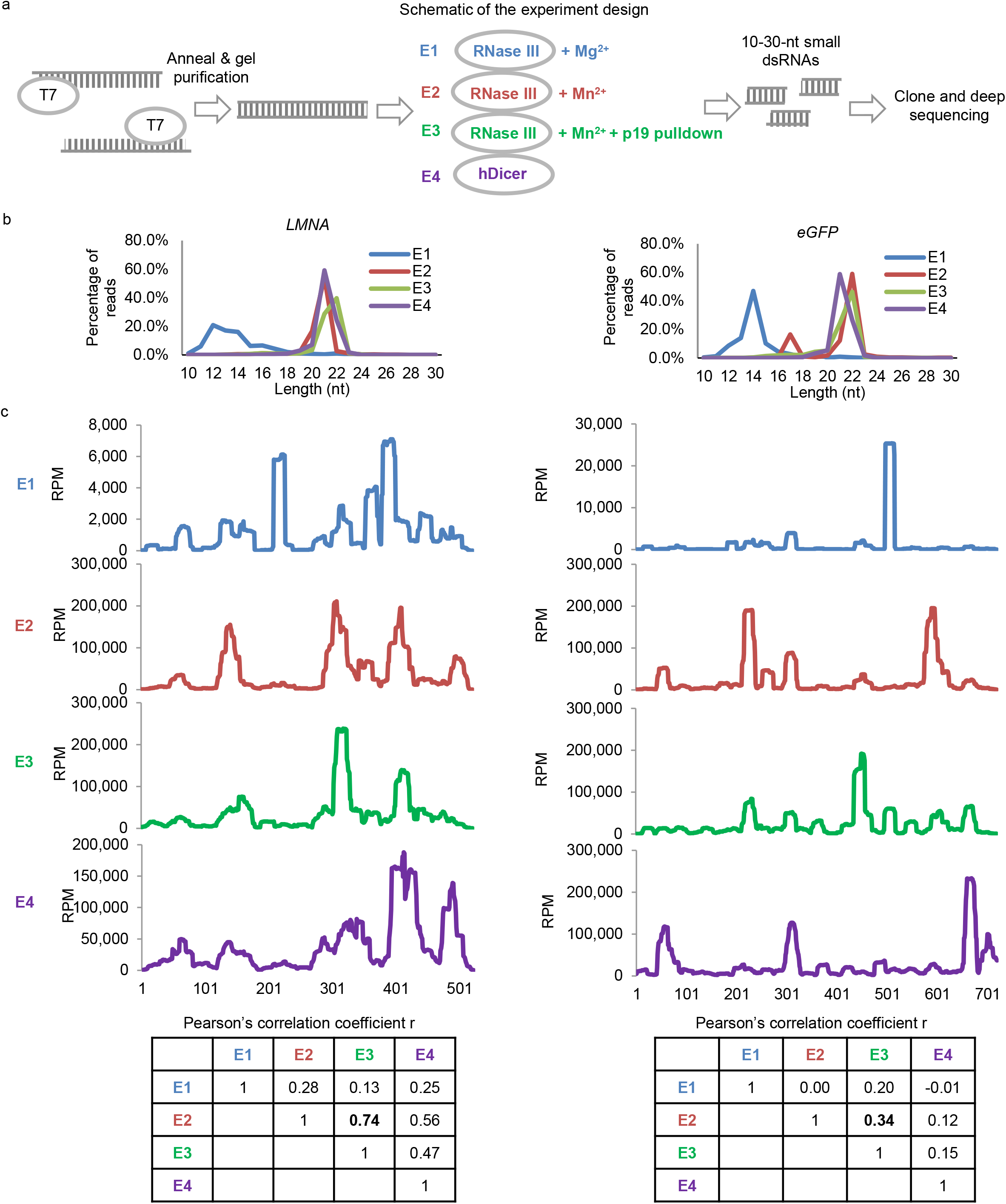
Hot spots of p19-captured dsRNAs are caused by RNase III, not p19 binding bias. a. Design of the experiment. b. Length distribution of small RNA deep sequencing data. c. Deep sequencing reads (combining sense and antisense reads) plotted along a 523 bp fragment of *LMNA* (left) and a 720 bp fragment of *eGFP.* RPM: reads per million. Bottom: pairwise correlation coefficient table calculated using small RNA sequence profiles.

**Supplementary Figure 3.**
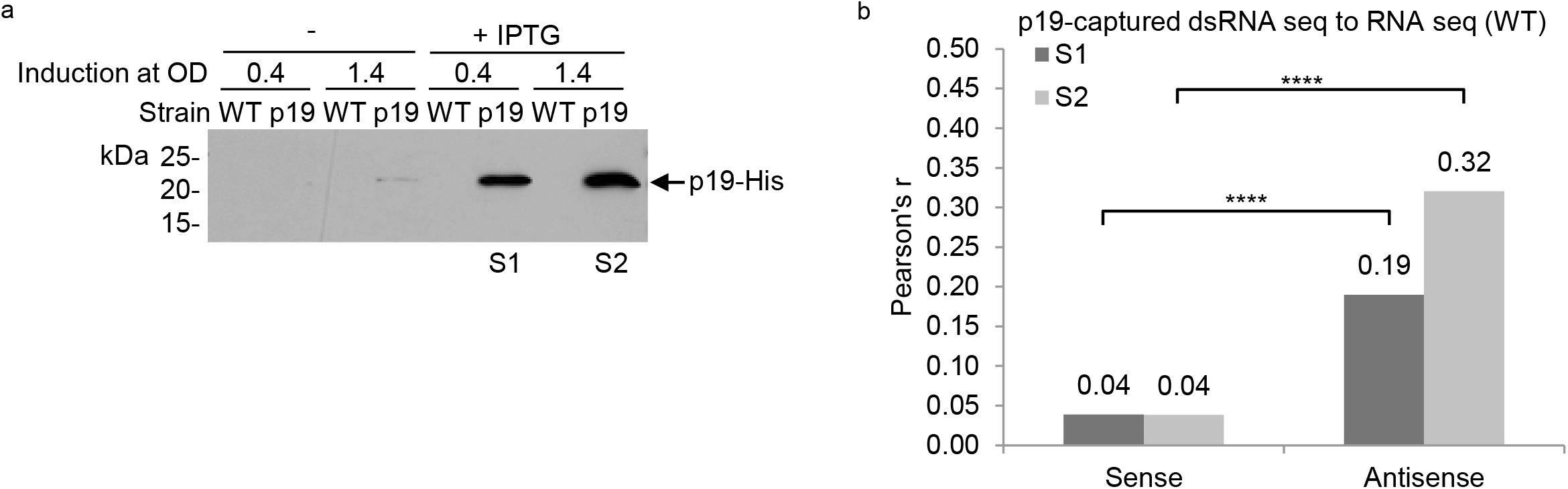
Features of p19-captured dsRNAs. a. Western blot probed with anti-His tag antibody. Samples were harvested from WT bacteria at S1 and S2 as indicated. b. Pearson’s correlation coefficient r, comparing p19-captured dsRNA (S1 orS2) and sense or antisense reads of total RNA sequencing data. ****: *p*-value<0.0001

**Supplementary Figure 4.**
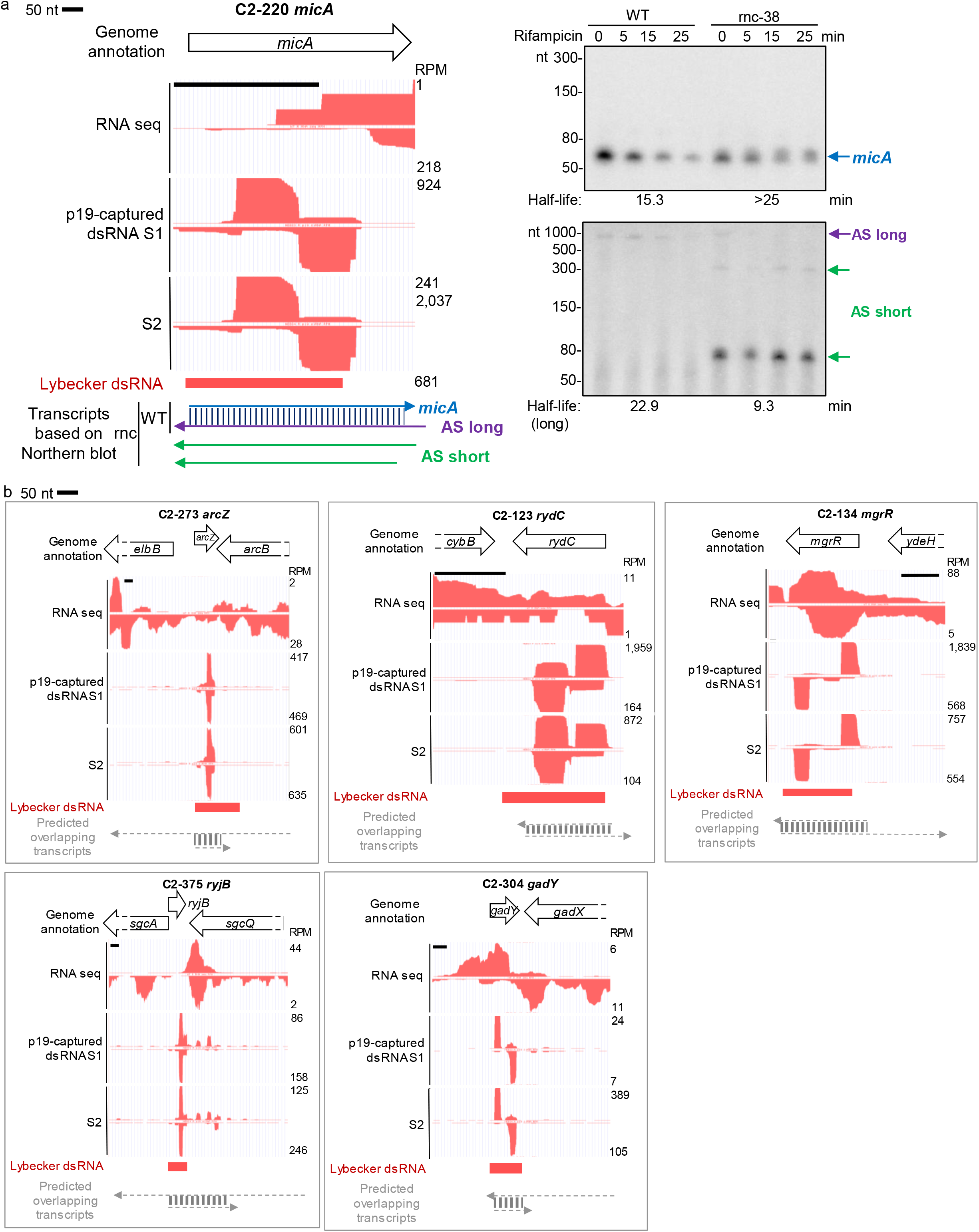
Examples of small RNA dsRNA clusters. a. *micA* locus and RNA half-life assay. Arrows indicate putative transcripts of small RNAs and their antisense transcripts. AS: antisense transcript. b. Other top 15 p19-captured dsRNA clusters involving small RNAs. Data plotted in UCSC genome browser as in Figure 3.

**Supplementary Figure 5.**
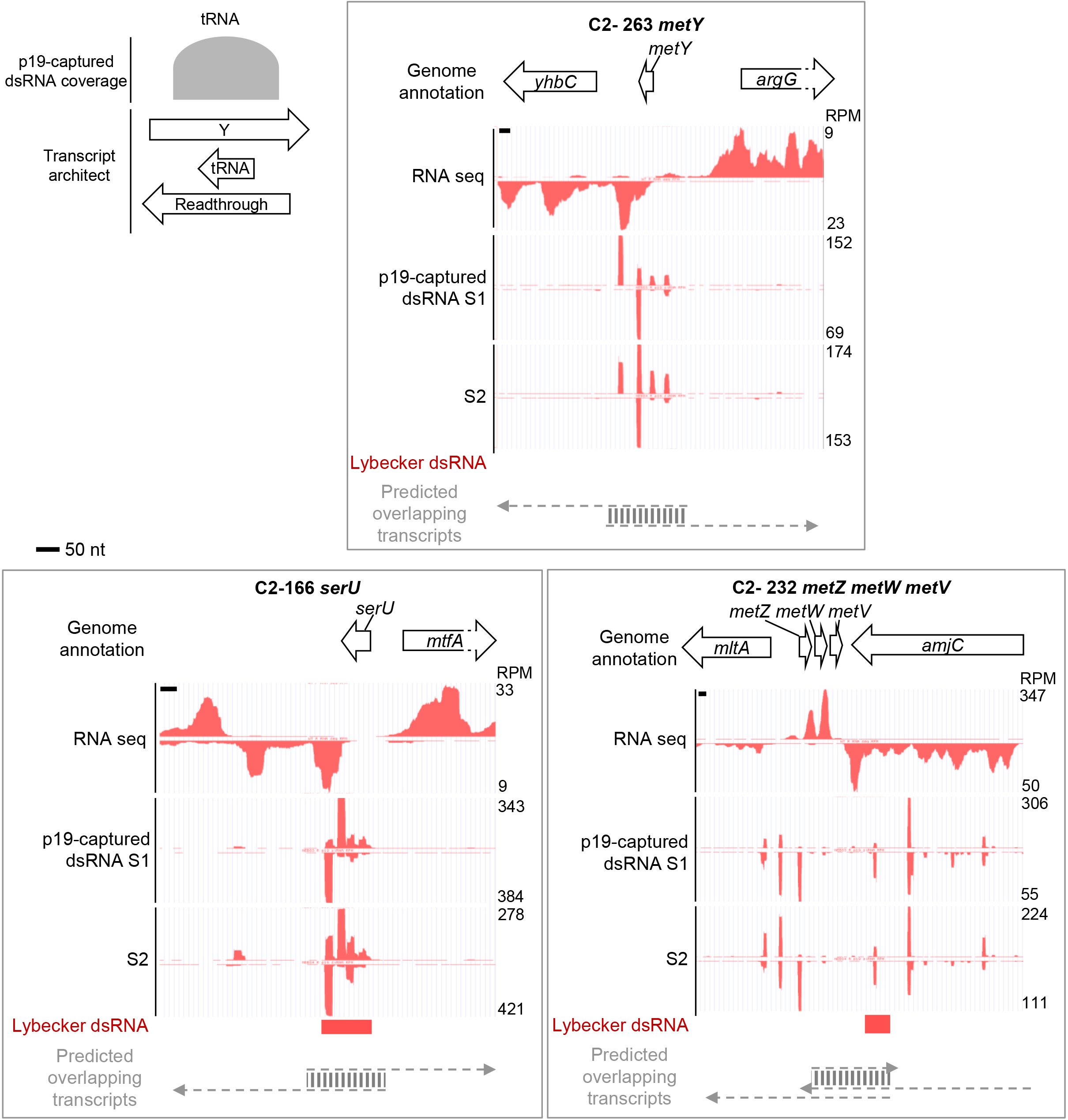
Examples of tRNA dsRNA clusters. Schematic showing overlapping antisense transcript for tRNA. Data plotted in UCSC genome browser as in Figure 3.

**Supplementary Figure 6.**
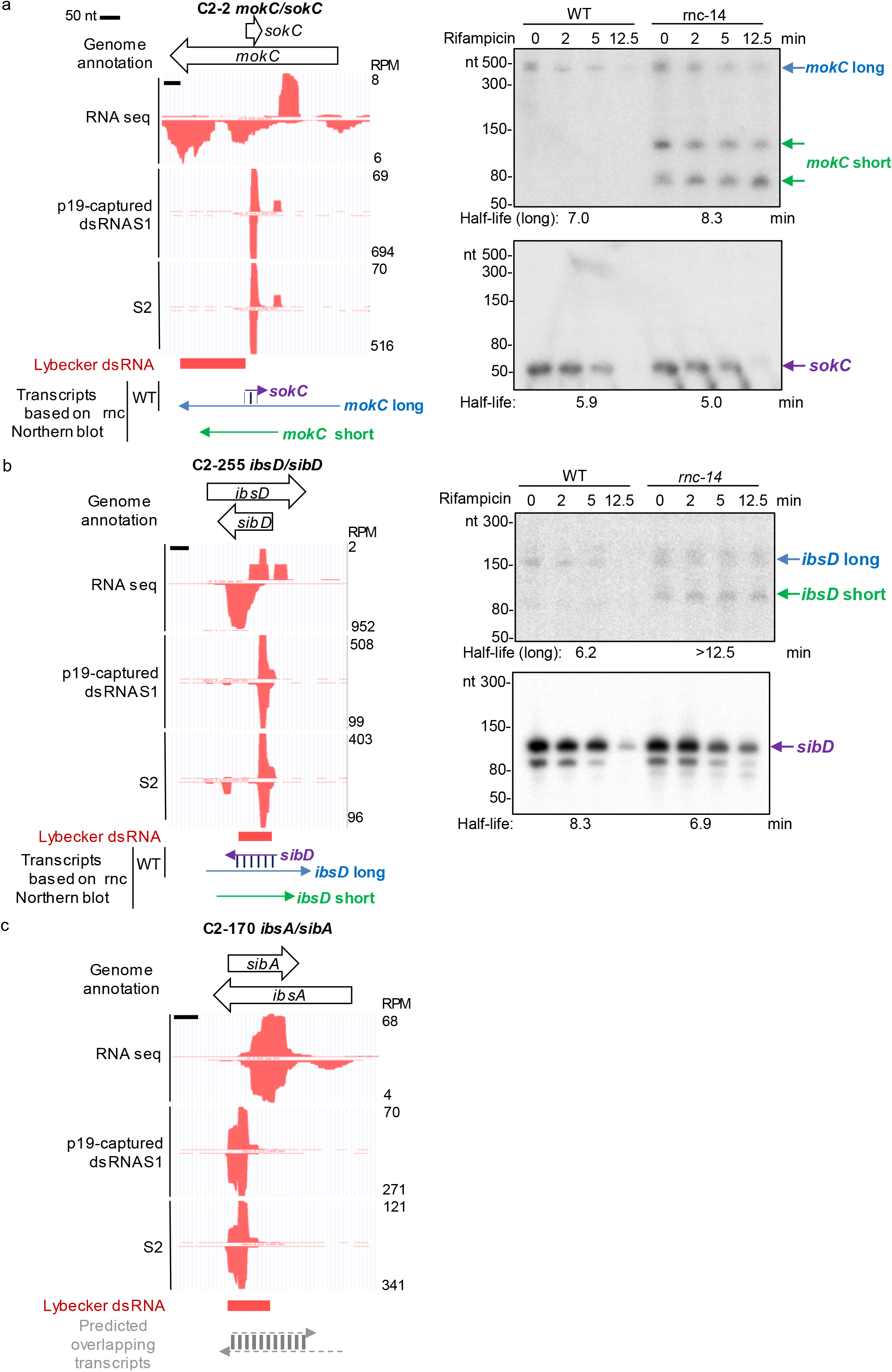
Type I TA loci generating dsRNAs. a. *mokC/sokC* locus and RNA half-life assay. b. *ibsD/sibD* locus and RNA half-life assay. c. *ibsA/sibA* locus. Arrows indicate putative RNA transcripts based on Northern blots. Data plotted in UCSC genome browser as in Figure 3.

**Supplementary Figure 7.**
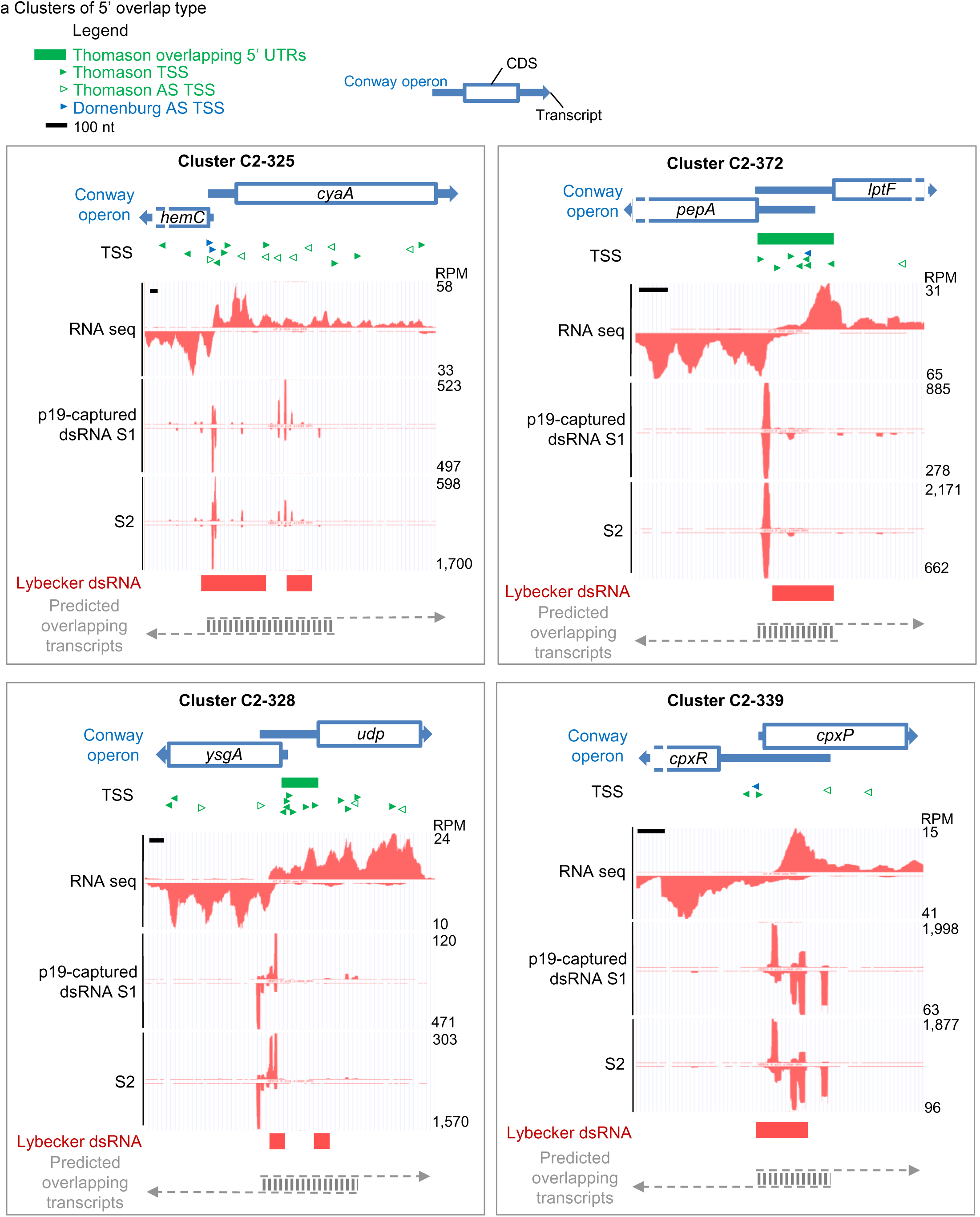
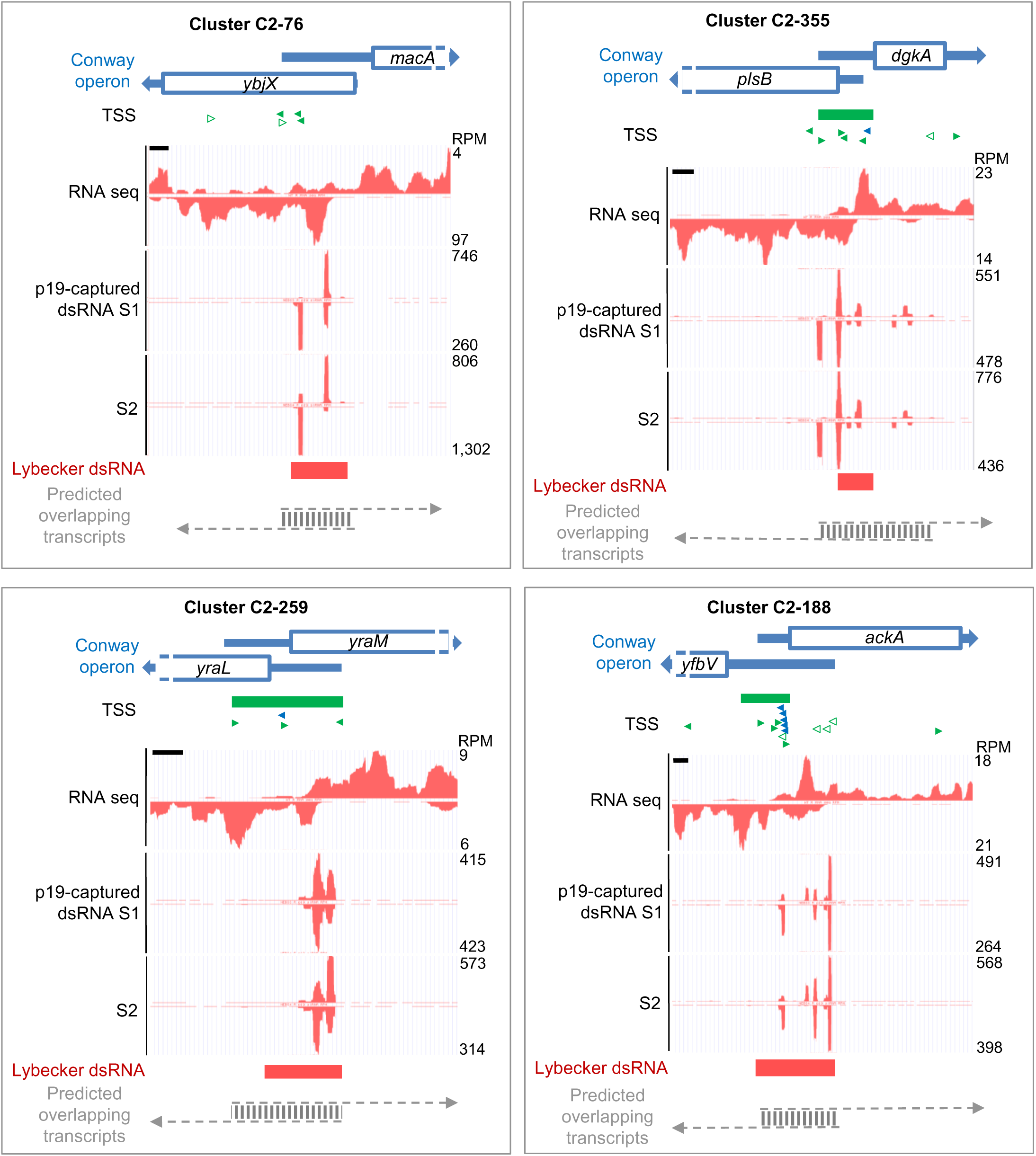
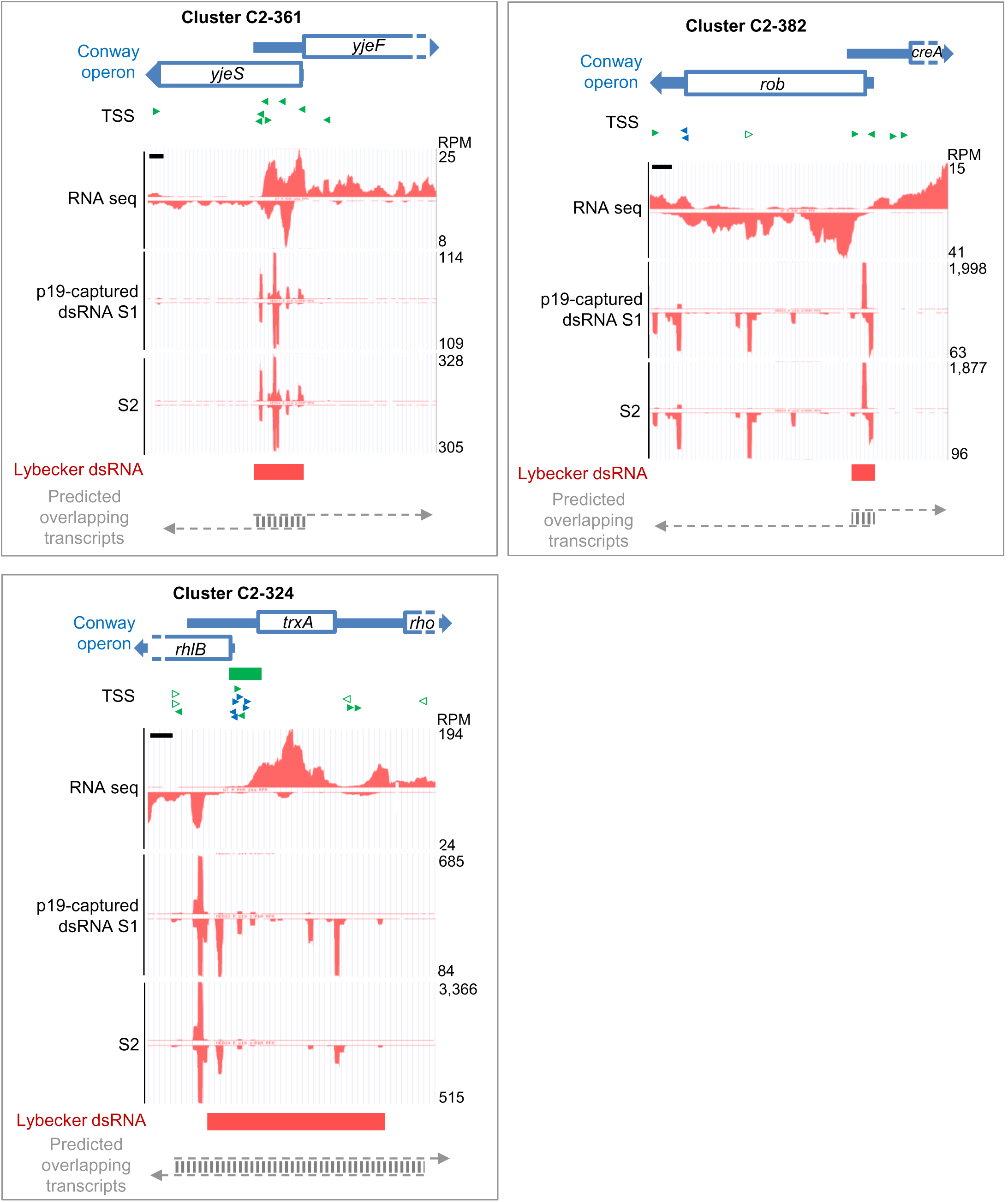
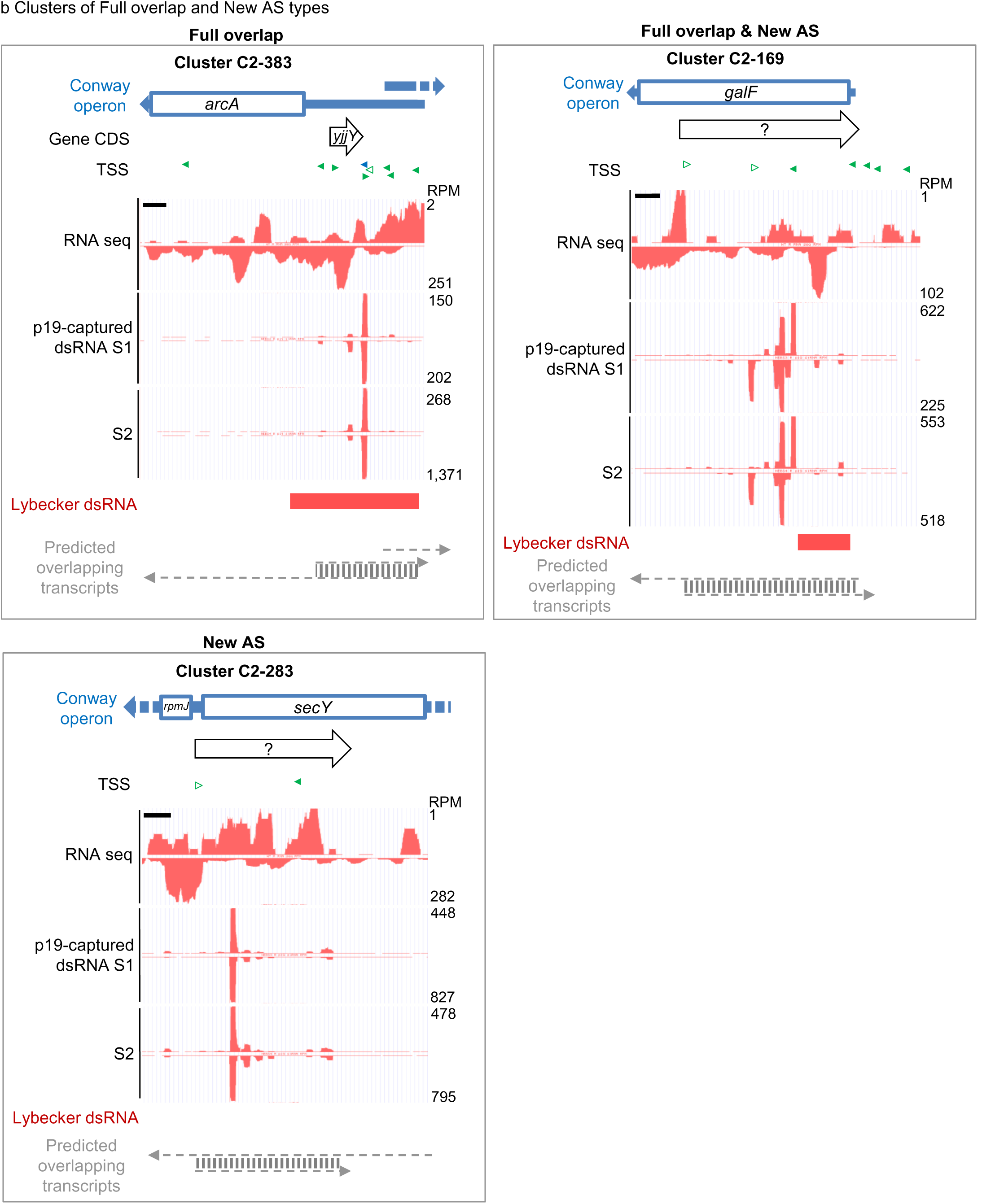
Other top 20 p19-captured dsRNA loci in coding genes. Data were plotted in the UCSC genome browser as in Figure 4. a. 5’ overlap type. b. Full overlap and New AS types.

**Supplementary Figure 8.**
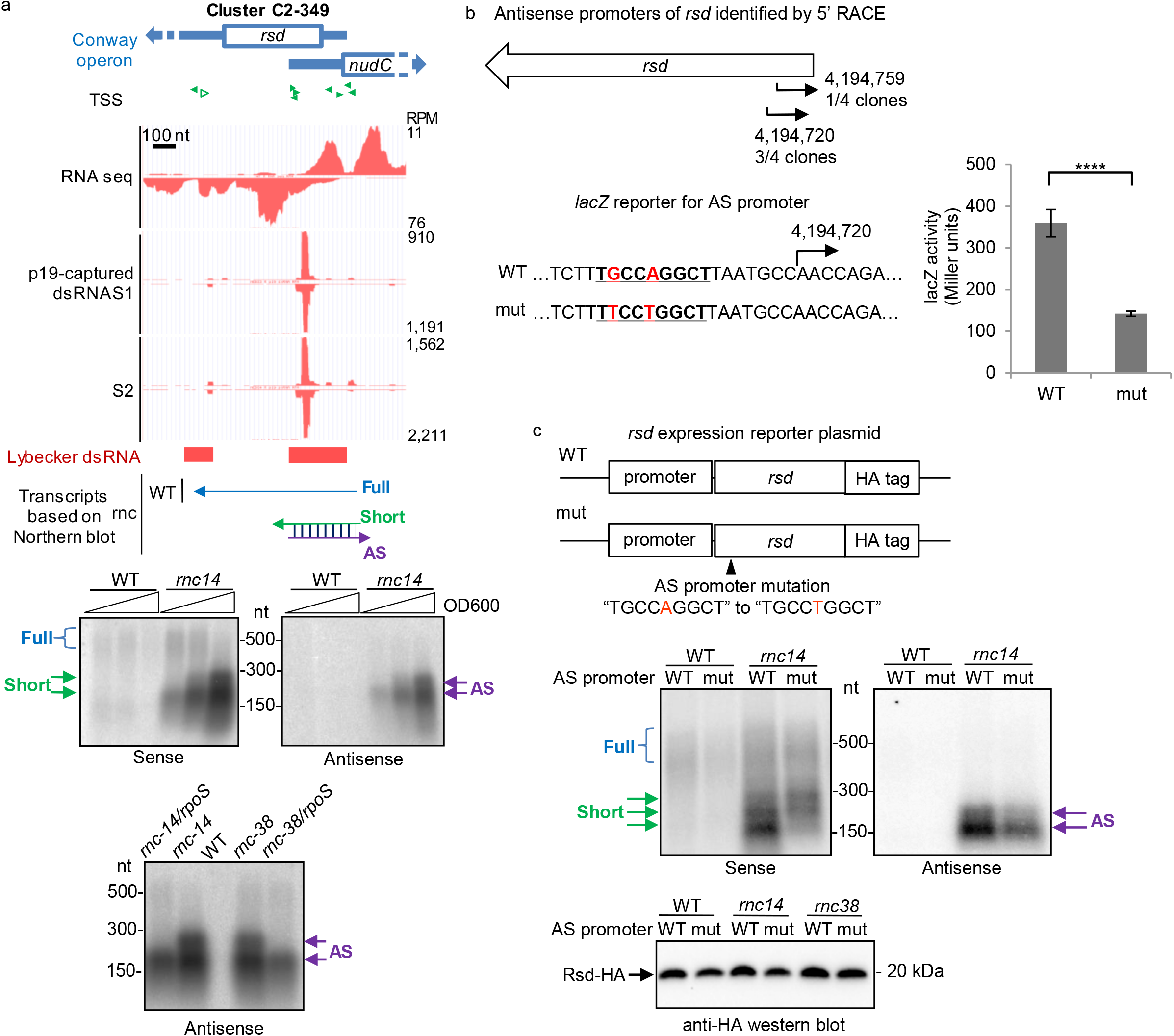
*rsd* antisense transcript. a. Top: p19-captured *rsd* dsRNA and RNA sequencing data plotted in UCSC genome browser and schematic of RNA transcript architecture of *rsd* locus; Middle: Northern blots probed for sense and antisense transcripts of *rsd;* Bottom: Northern blots for *rsd* antisense transcript in WT, *rnc* mutants, and *rnc/rpoS* double mutants. Arrows indicate detected putative RNA transcripts with color scheme matching the schematic. b. Cloning of putative *rsd* antisense transcript start site by 5’ RACE. Promoter activity of predicted antisense promoter and its mutation using a *lacZ* reporter. An extended −10 rsd antisense promoter element is underlined. ****: *p*-value<0.0001. c. *rsd* expression from WT and mutant antisense promotors in *E. coli* using a plasmid system. Top: schematic of the *rsd* expression reporter plasmids with or without antisense promoter mutation; Middle: Northern blot probed for sense and antisense transcripts of *rsd;* Bottom: Western blot to detect HA-tagged Rsd protein in WT and *rnc* mutants. AS: antisense transcript.

**Supplementary Figure 9.**
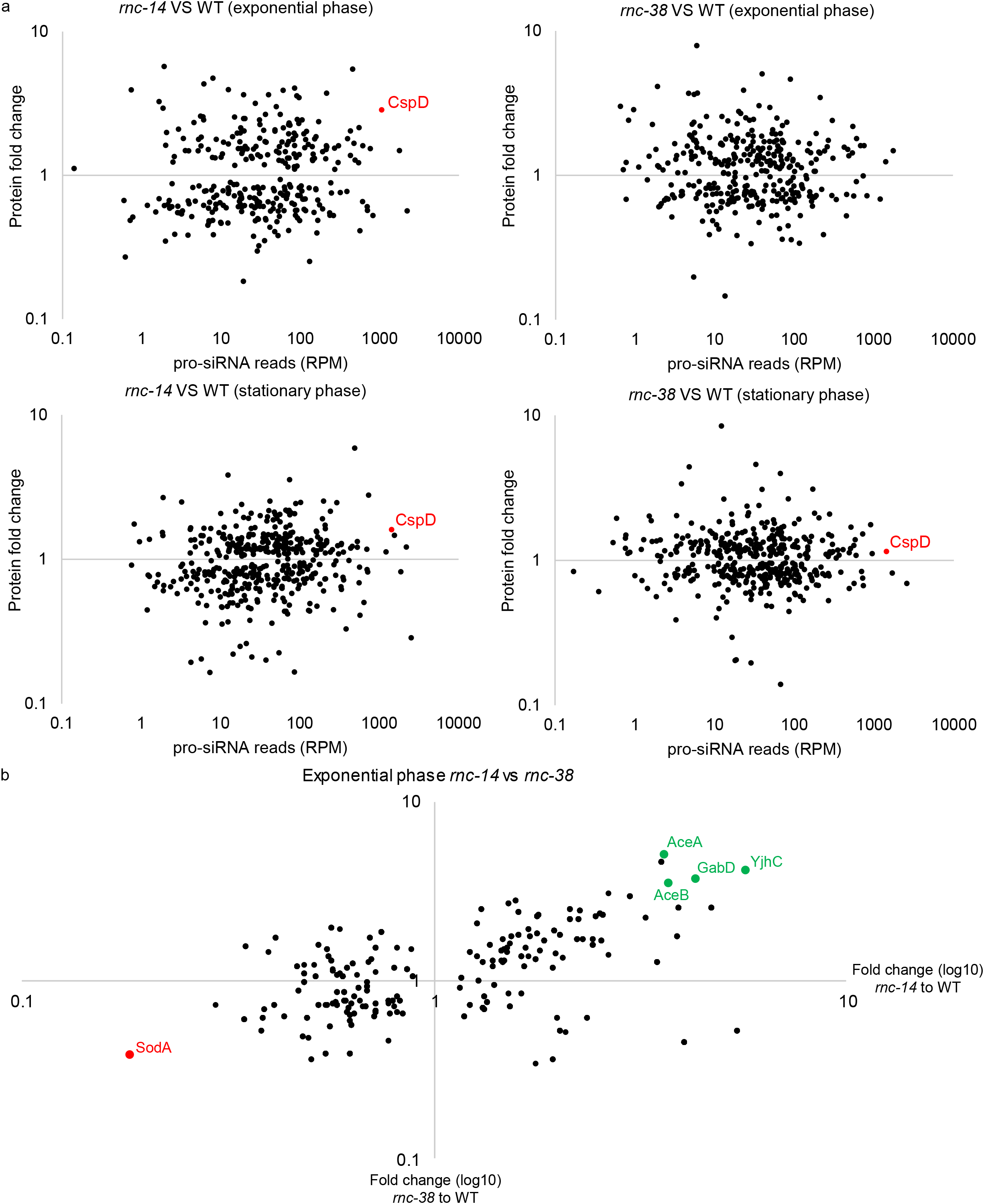
Quantitative proteomics comparing *E. coli rnc* mutants with WT. a. The protein fold change *(rnc* to WT) was plotted against the p19-captured dsRNA abundance of the gene. b. XY plot for protein fold change in *rnc-14* (X) and *rnc-38* (Y). Genes that are consistently changed in both *rnc* mutants are marked (upregulated genes in green and downregulated gene in red).

**Supplementary Figure 10.**
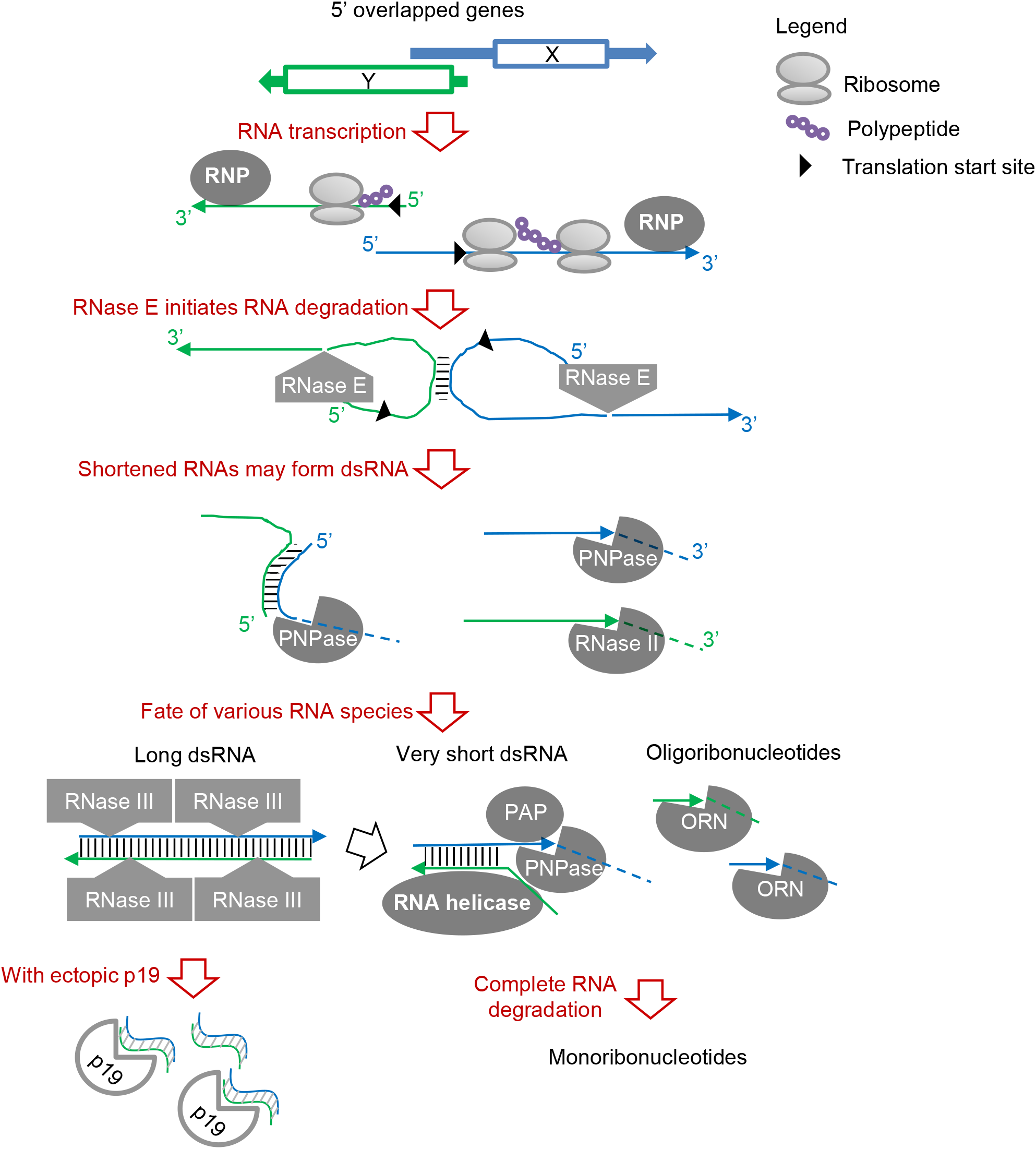
Model for RNA decay in *E. coli.* RNP: RNA polymerase; PAP: Poly(A) polymerase; PNPase: Polyribonucleotide nucleotidyltransferase; ORN: Oligoribonuclease.

**Supplementary Table 5.** Major studies on identifying RNase III targets in genome-wide scale in bacteria.

|  | **Lasa et al. PNAS,**<br>**2011 (6)** | **Lioliou et al. PLOS**<br>**Genetics 2012 (7)** | **Lybecker**<br>**et al.**<br>**PNAS, 2013 (8)** | **Le Rhun et al. NAR,**<br>**2017 (9)**<br>**Gordon et al. mBio,**<br>**2017 (10)**<br>**Altuvia et al. NAR,**<br>**2018 (11)** | **This work** |
| --- | --- | --- | --- | --- | --- |
| **Method** | Deep sequencing of<br>very short RNAs | dsRNA pulldown by<br>RNase III catalytic<br>mutant | dsRNA pulldown by<br>anti-dsRNA J2<br>antibody in *rnc*<br>mutant | Special deep sequencing<br>methods and advanced<br>analysis comparing WT<br>and *rnc* mutant to<br>identify ends of RNase<br>III cleaved RNAs | Ectopic expression of<br>*Tombusvirus* p19 to<br>capture RNase III<br>products in cells |
| **Tested bacteria** | *S. aureus*, *E. faecalis*,<br>*L. monocytogenes*, *B.*<br>*subtilis*, *S. Enteritidis* | *S. aureus* | *E. coli* | *E. coli*, *S. pyogenes* | *E. coli* |
| **Require *rnc* mutant** | Yes | No | Yes | Yes | No |
| **Precise RNase III**<br>**cleavage site** | No | No | No | Yes | Yes |

**Supplementary Table 6.**
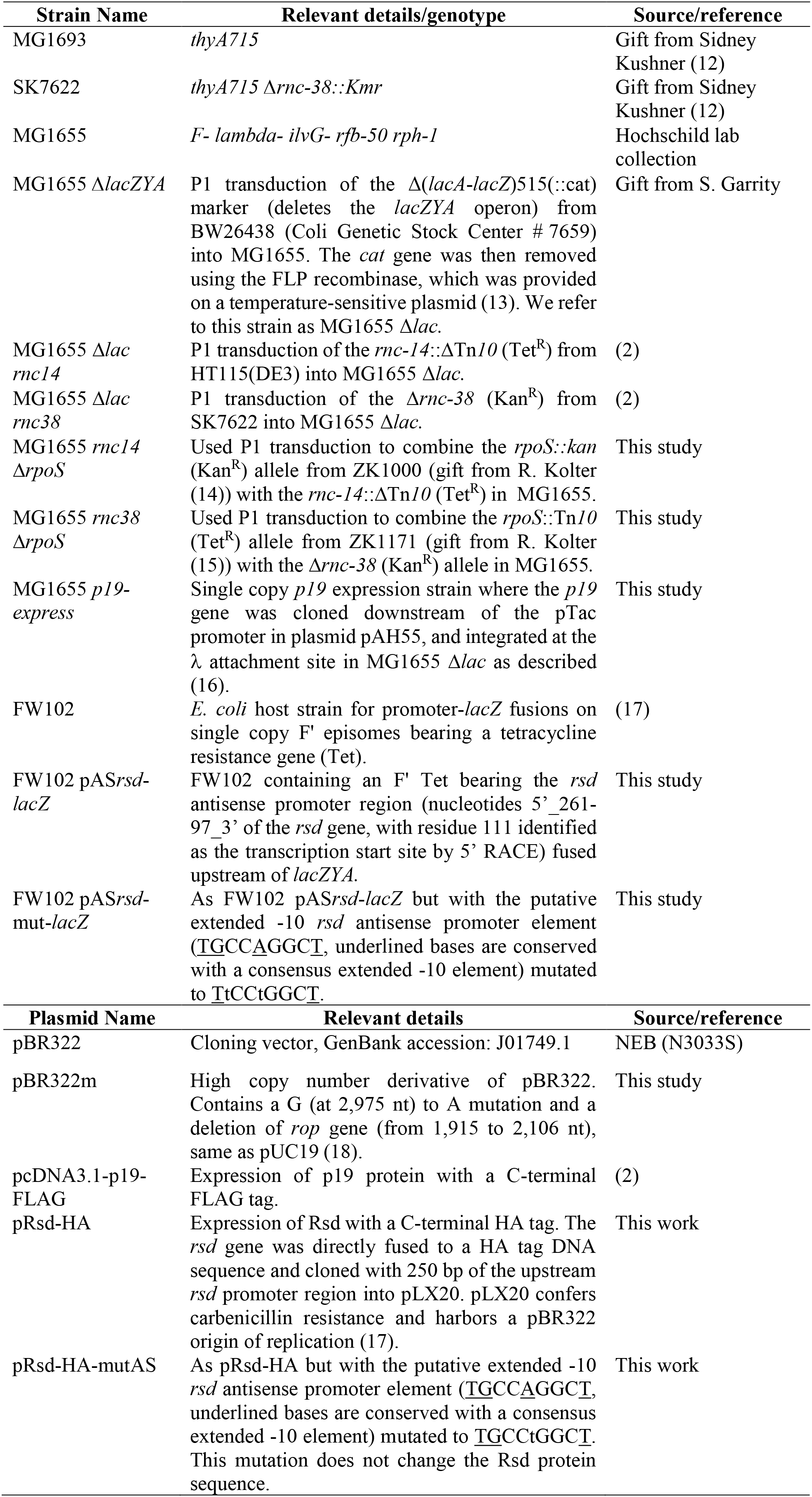
List of *E. coli* strains and plasmids used in this study.

**Supplementary Table 7.** Summary of sample information and sequence alignment results for deep sequencing experiments.

| **NCBI BioSample**<br>**accession** | **Type** | **Sample name** | **Total reads** | **Aligned reads** | **Percentage**<br>**aligned** |
| --- | --- | --- | --- | --- | --- |
| SAMN10659748 | Small RNA | p19-captured dsRNA S1 | 85,262,844 | 22,121,763 | 25.9% |
| SAMN10659749 | Small RNA | p19-captured dsRNA S2 | 87,388,195 | 30,995,122 | 35.5% |
| SAMN10659750 | Small RNA | p19-captured dsRNA pcDNA3.1-p19-FLAG from MG1693 | 780,247 | 546,389 | 70.0% |
| SAMN10659751 | Small RNA | p19-captured dsRNA pcDNA3.1-p19-FLAG from SK7622 | 64,854 | 42,947 | 66.2% |
| SAMN10659752 | Total RNA | Total RNA WT | 11,868,604 | 10726837 | 90.4% |
| SAMN10659753 | Total RNA | Total RNA rnc-14 | 24,260,091 | 21,758,743 | 89.7% |
| SAMN10659754 | Total RNA | Total RNA rnc-38 | 8,022,580 | 7,896,135 | 98.4% |
| SAMN10659756 | Small RNA | E1 LMNA | 1,356,245 | 1,117,580 | 82.4% |
| SAMN10659757 | Small RNA | E2 LMNA | 878,954 | 813,092 | 92.5% |
| SAMN10659758 | Small RNA | E3 LMNA | 781,594 | 665,761 | 85.2% |
| SAMN10659759 | Small RNA | E4 LMNA | 6,522,249 | 6,175,984 | 94.7% |
| SAMN10659760 | Small RNA | E1 eGFP | 8,750,398 | 7,922,440 | 90.5% |
| SAMN10659761 | Small RNA | E2 eGFP | 7,247,371 | 5,819,972 | 80.3% |
| SAMN10659763 | Small RNA | E3 eGFP | 4,715,011 | 3,555,925 | 75.4% |
| SAMN10659764 | Small RNA | E4 eGFP | 1,660,915 | 1,549,318 | 93.3% |

